# SeqScreen: Accurate and Sensitive Functional Screening of Pathogenic Sequences via Ensemble Learning

**DOI:** 10.1101/2021.05.02.442344

**Authors:** Advait Balaji, Bryce Kille, Anthony D. Kappell, Gene D. Godbold, Madeline Diep, R. A. Leo Elworth, Zhiqin Qian, Dreycey Albin, Daniel J. Nasko, Nidhi Shah, Mihai Pop, Santiago Segarra, Krista L. Ternus, Todd J. Treangen

## Abstract

The COVID-19 pandemic has emphasized the importance of detecting known and emerging pathogens from clinical and environmental samples. However, robust characterization of pathogenic sequences remains an open challenge. To this end, we developed SeqScreen, which can accurately characterize short nucleotide sequences using taxonomic and functional labels, and a customized set of curated Functions of Sequences of Concern (FunSoCs) specific to microbial pathogenesis. We show our ensemble machine learning model can label protein-coding sequences with FunSoCs with high recall and precision. SeqScreen is a step towards a novel paradigm of functionally informed pathogen characterization and is available for download at: www.gitlab.com/treangenlab/seqscreen

## Introduction

Rapid advancements in synthesis and sequencing of genomic sequences and nucleic acids have ushered in a new era of synthetic biology and large-scale genomics. While the democratization of reading and writing DNA has greatly enhanced our understanding of large-scale biological processes[1], it has also introduced new challenges[2]. Robust characterization of genetically engineered or *de novo* synthesized pathogens has never been more relevant, and the importance of detecting and tracking naturally evolving and emerging pathogenic sequences from the environment cannot be overstated. Open challenges that represent barriers to accurate detection include, but are not limited to, (i) the role of abiotic and environmental stress response genes in virulence, (ii) the presence of seemingly pathogenic sequences in commensals, (iii) host-specific pathogen virulence, and (iv) interplay of different genes to generate pathology[3]. Accurate and sensitive detection of pathogenic markers has also been confounded by the difficulty of characterizing multifactorial microbial virulence factors in the context of the biology of the host[4]. The limited number of publicly available databases to annotate and identify specific pathogenic elements within sequencing datasets further exacerbates the problem. Furthermore, due to difficulties with automated annotations and the lag between experimental results and sequence annotations, identifying sequences involved in pathogenesis is an ongoing challenge[5,6]. Gene Ontology (GO) terms were not designed to solely capture the nuanced biological processes and molecular functions specific to pathogens, and the pathogenesis GO term (GO:0009405), which labels >275K UniProt accessions, was recently made obsolete, with the final notice given in March 2021 (https://github.com/geneontology/go-annotation/issues/3452). Thus, there exists an urgent need in the community for a tool that can accurately characterize genomic sequences in the context of functional pathogen detection and identification, thereby sensitively capturing sequences of concern (SoCs) in each sample[3]. With respect to computational approaches for pathogen characterization, much recent progress has been made specific to taxonomic classification from isolates and metagenomic datasets. For example, probabilistic models leveraging k-mer genotyping and logistic regression analysis to identify k-mers indicative of antibiotic resistance have shown promise [7]. Other tools incorporating statistical frameworks for predicting markers of pathogenicity from sequencing data include PathoScope[8,9] and SURPI[10]. The former utilizes sequence quality and mapping quality as parts of a Bayesian model to rapidly compute posterior probabilities of matches against a database of known biological agents, while the latter uses either Scalable Nucleotide Alignment Program (SNAP)[11] based alignments to bacterial or viral databases and in some cases RAPSearch[12] for more sensitive identification. Both tools also had separate releases, Clinical PathoScope[13] and SURPI+[14], specifically focused on pathogen characterization from clinical samples. Another k-mer based tool by CosmosID[15], precomputes reference databases (reference genomes as well as virulence and antimicrobial resistance markers) to create a phylogeny tree of microbes as well as variable-length k-mer fingerprint sets for each branch and leaf of the tree. Sequencing reads are then scanned against these unique fingerprint sets for detection and taxonomic classification. The statistics derived are then refined using predefined internal thresholds and statistical scores to exclude false positives and fine grain taxonomic and relative abundance estimates. Evaluations of this approach have shown that CosmosID achieves a high level of sensitivity in antibiotic resistance gene detection for predicting staphylococcal antibacterial susceptibility[16]; however, this is not an open-source tool and was not further evaluated in this study.

However, despite recent progress, all of the aforementioned methods either: i) assume the presence of the entire genome, ii) ignore functional information, or iii) are ill-equipped to analyze individual short sequence lengths typical of synthesized oligonucleotides. Previous benchmarking studies on microbial identification from metagenomes have shown that there exists a crucial tradeoff between taxonomic resolution and accuracy given the current state-of-the-art tools[17]. Furthermore, taxonomic id is often a poor proxy for pathogenicity. While modern computational methods have tackled aspects of this problem by focusing of various types of pathogenic markers, there exists a gap in computational tools and annotation frameworks able to accurately identify known and emerging pathogens from environmental samples [18]. It is precisely this gap that we aimed to fill with SeqScreen. Previously, we introduced a proof-of-concept framework[19] for robust taxonomic and functional characterization of nucleotide sequences of interest. Here, we build upon the earlier framework and present a robust and comprehensive tool based on ensemble machine learning and functions of sequences of concern (FunSoCs) for pathogen identification and detection. Our system, SeqScreen, combines alignment-based tools, ensemble machine learning classifiers, curated databases, and novel curation-based labelling of protein sequences with pathogenic functions, to identify sequences of concern in high throughput sequencing data. Through careful, manual assignment of pathogenic functions based on published investigations of each sequence, SeqScreen depends on high quality training data to predict FunSoCs accurately. The SeqScreen FunSoC database has been pre-computed with our ensemble machine learning classifiers, so the SeqScreen software does not train the machine learning classifier or run machine learning in real time, making the analysis more streamlined and the results consistently reproducible and reviewable. SeqScreen aspires to be the first tool to combine human interpretability and machine learning-based classification in a human-in-the-loop construct to provide a holistic solution towards classifying pathogens and offers a novel functional framework for pathogen identification in contrast to existing tools.

## Results

### Comparison of FunSoCs to previous pathogen detection frameworks

Previous pathogen detection methods have mainly relied on the Virulence Factor Database (VFDB) as a training and validation dataset to detect markers of pathogenicity from Next Generation Sequencing (NGS) data[20,21]. VFDB contains a set of more than 3400 core sequences that aim to capture Virulence Factors (VFs) from 30 different genera of medically relevant bacterial pathogens[22]. There have been five updates describing VFDB since the original announcement published in 2005, with the latest being in 2019[22–26]. A close inspection of VFDB sequences revealed some limitations to basing our tool on this framework. There is little rationale provided for why these sequences and not others are included. No Gene Ontology terms, or other functional annotations, are used to describe individual sequences. VFDB contains many proteins that contribute to flagellar production. Flagellar components are recognized by pattern recognition receptors of the innate immune system and can thus precipitate an inflammatory reaction, but they are found in both pathogenic and non-pathogenic species. In any case, the flagellar synthases could only remotely be considered pathogenic. A vast majority of the sequences also were involved in secretomes or general secretion pathways and did not fully capture the diversity of VFs. To address these limitations, our curation team developed the FunSoC framework that improved upon the functional inclusion criteria and developed a new set of proteins consisting of 1433 training sequences that contained different GO terms that represented the underlying FunSoCs with each sequence having at least one FunSoC annotation. **Fig. 1 A** shows the overlap between the distinct GO terms from the VFDB core sequences, and the training set used in our study. The SeqScreen training dataset contained 12086 GO terms compared to just 657 retrieved from the VFDB sequences. The lack of functional information in VFDB was also observed by comparing the annotation scores (**Fig. 1 B**) of sequences as specified in UniProt. The annotation score of VFDB core sequences was overwhelmingly 1 (out of 5), whereas the sequences in our training dataset were carefully curated to include proteins that had annotation scores above 3 with a median score of 4, indicating a higher degree of confidence in its functional annotation within UniProt. Hence, the FunSoCs offered a high-quality, manually curated training dataset consisting of proteins with a wider variety of functional annotations that underly different mechanisms of microbial pathogenicity.

**Fig. 1.**
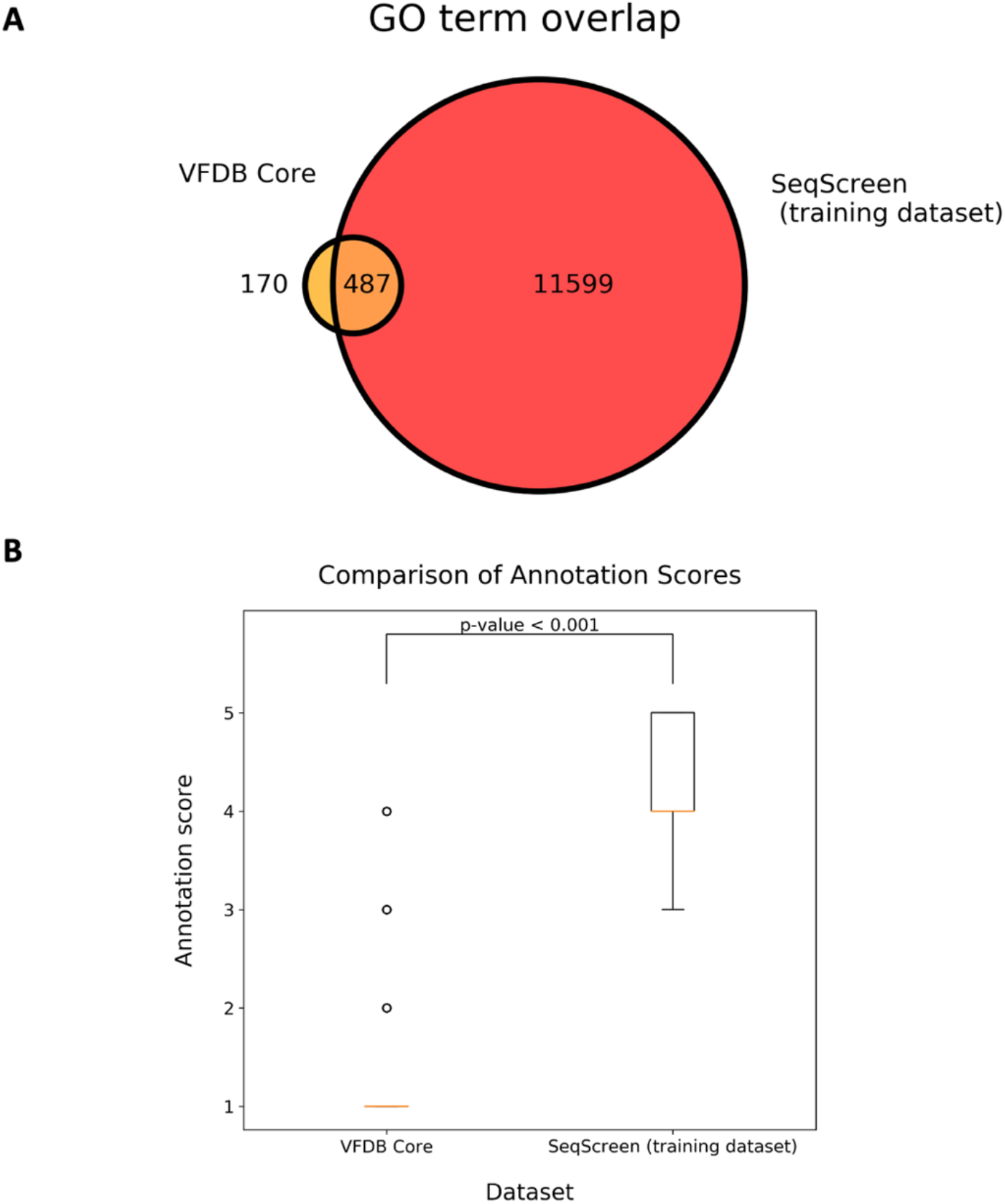
Comparison of VFDB to SeqScreen Biocurator Database. A. Venn-diagram shows the number of GO terms captured by VFDB Core sequences, the SeqScreen training dataset labelled by biocurators, and their overlap. B. Box-plot showing the comparison of annotation scores (1-5) of the associated UniProt/UniParc IDs between VFDB Core sequences and SeqScreen training data. The p-value was calculated using the Mann-Whitney U test.

### Pipeline Overview and Module Descriptions

The SeqScreen pipeline was built using Nextflow[27], a domain-specific language for creating scalable and portable workflows. SeqScreen combines various stages in separate Nextflow modules and is available as an open-source tool on bioconda (https://anaconda.org/bioconda/seqscreen**). Fig. 2** illustrates the various modules and five main workflows in SeqScreen. SeqScreen can be run in two different modes -*default* (i.e., fast) mode and *-sensitive* mode. The default *fast* mode runs a limited set of pipelines that are tuned to rapidly annotate sequences in an efficient performance-centric approach. The *sensitive* mode (using the *--sensitive* flag) uses much more accurate and sensitive BLASTN-based alignments[28] and outlier detection[29] steps for taxonomic characterization. Further, for sensitive functional annotations it uses BLASTX to identify hits to the curated UniRef100 database. The modular nature of the pipeline offers advantages in terms of ease of updating or replacing specific software modules in the future versions if new bioinformatics tools and databases are shown to outperform its current modules and workflows. SeqScreen accepts nucleotide FASTA files as input, assuming one protein-coding sequence is present within each query sequence of the FASTA file. Each input file is verified for the correctness of the FASTA format and then passed on to the initialization workflow in *sensitive* mode, which first converts ambiguous nucleotides to their corresponding unambiguous options and performs six-frame translations of nucleotide to amino acid sequences for input into downstream modules like RAPSearch2[12], which accepts amino acid sequence as input. After initialization, the sequences pass through various downstream modules that add taxonomic and functional annotations to the sequences that inform its FunSoC assignment. The downstream modules depend on the mode the user runs SeqScreen in; *-default* (DIAMOND[30] and Centrifuge[31]) or *-sensitive* (BLASTX, MUMmer[32]+REBASE [33]and MEGARes[34]). FunSoC assignment of query sequences is carried out by transferring the FunSoC labels of the target proteins in our database identified during functional annotation. This database containing mappings from individual UniProt Ids to FunSoCs to is precomputed from the predictions of the ensemble machine learning classifier. Training data for the classifier was obtained from manual curations of literature and databases by our team of expert biocurators. The precomputed FunSoC database obviates the need to run the classifier in real-time thereby increasing the efficiency of the SeqScreen pipeline. All analyses in this study were performed with SeqScreen *-default* mode, other than the SeqMapper-focused analysis that was run in sensitive mode. Each of the individual workflows of the SeqScreen pipeline are discussed in more detail below.

**Fig. 2.**
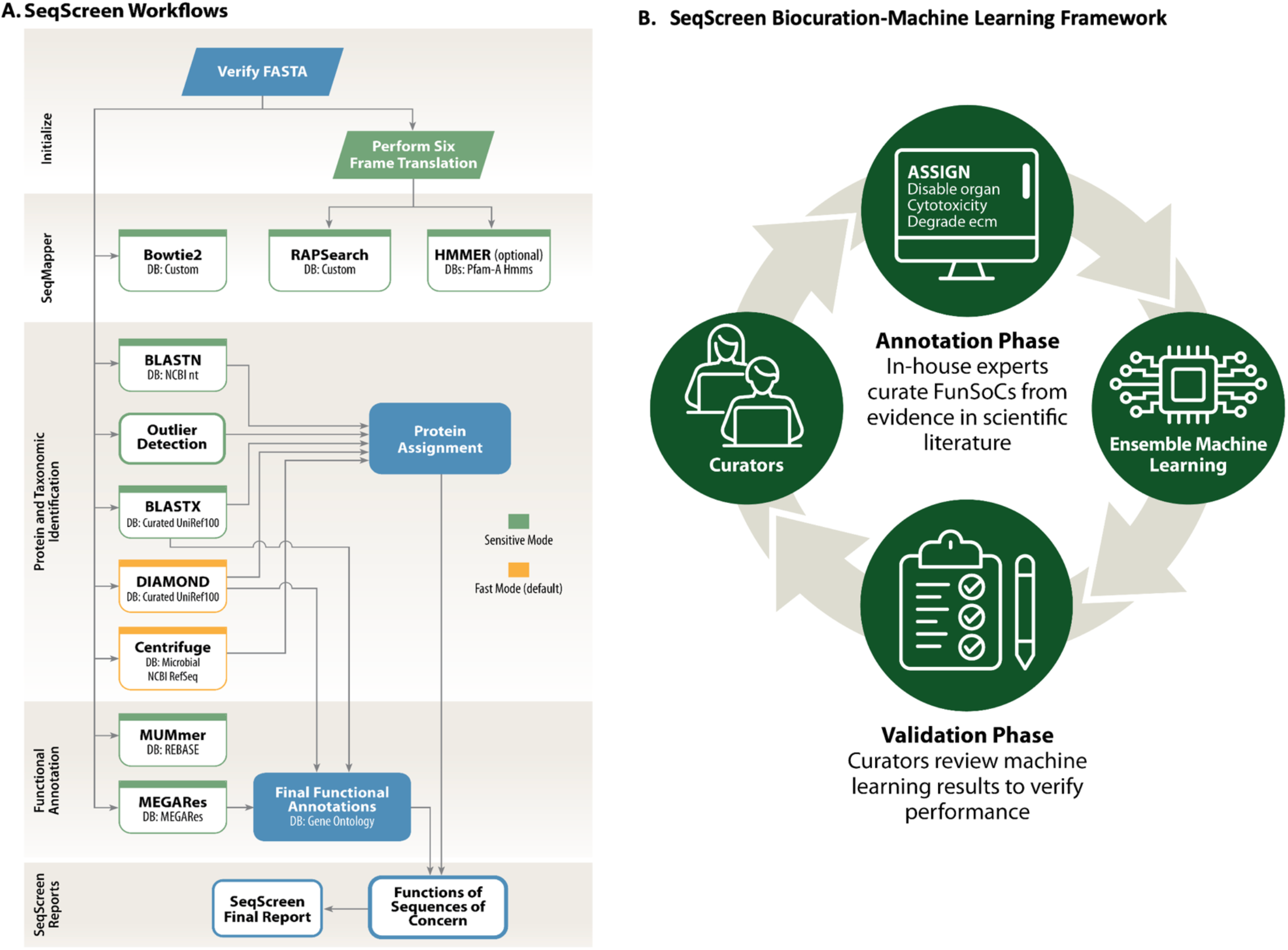
SeqScreen overview. (A.) SeqScreen Workflow: This figure outlines the various modules and workflows of the SeqScreen pipeline. Boxes in green indicate that these modules are only run in the sensitive mode. The boxes in yellow are run in the fast mode, while the ones in blue are common to both modes. In addition to the two different modes, SeqScreen also contains optional modules that can be run based on the parameters provided by the user. (B.) SeqScreen Human-in-the-loop Framework: Includes initial annotation and curation of training data by manual curation. The data is used to train Ensemble ML models. The results obtained and selected feature weights are passed on back to biocurators to fine tune features and uniport queries which form a new set of refined training data for the Ensemble model.

### SeqScreen workflow #1: Initialize

Each run is initialized by first checking the input fasta file and verifying it to be error free. Some common errors that are screened for include headers with empty sequences, duplicate headers, and invalid or ambiguous bases. SeqScreen also checks for suspiciously long sequences depending on a user-controlled parameter (--max_seq_size). In addition to quality control for input sequences, the sensitive mode also contains the six-frame translation module to convert the nucleotide sequence into amino acids to input to the SeqMapper module.

### SeqScreen workflow #2: SeqMapper

The SeqMapper workflow is part of the *sensitive* mode of SeqScreen and includes additional features, such as detecting Biological Select Agents and Toxins (BSAT) sequences through efficient sequence alignment methods. We use a two-pronged approach by analyzing both the nucleotide and amino acid sequence alignments to BSAT reference genomes using Bowtie2[35] and RAPSearch2[12], respectively. While this workflow is only limited to reporting hits to BSAT genes and proteins, downstream workflows are used to capture and collate whether a gene is of interest at a functional level (e.g., functional differentiation between BSAT housekeeping and toxin hits are not delineated at this step). This workflow is sensitive to detect BSAT sequences, but not precise in differentiating BSAT sequences from their near neighbors. The BSAT sequences were primarily derived from the following website: https://www.selectagents.gov/sat/list.htm and the full contents of the BSAT Bowtie2 database is available at: https://rice.box.com/s/6c5xl0qcu66xbuf3n8yp4fkf9cfwn3wv. In addition to the above databases, users can also optionally obtain other features of interest, such as HMMs identified by HMMER[36] from Pfam[37] proteins by using the optional HMMER module in SeqScreen.

### SeqScreen workflow #3: Protein and Taxonomic Identification

In the taxonomic classification workflows for both *fast* and *sensitive* modes we rely on widely used state-of-the-art alignment-based tools to classify sequences. SeqScreen obtains alignments to both DNA and amino acid databases. While aligning to amino acid databases provides taxonomic information as well as functional information, aligning to nucleotide databases provides additional sensitivity, especially for non-coding regions. The taxonomic classification module for *fast* mode is an ensemble of DIAMOND and Centrifuge, two established and widely used tools for protein alignment and taxonomic classification. First, DIAMOND is used to align the input sequences to a reduced version of the UniRef100 database. DIAMOND is an open-source software that is designed for aligning short sequence reads and performs at approximately 20,000 times the speed of BLASTX with similar sensitivity. Our reduced version of the UniRef100 database[38] only contains proteins with a high annotation score. Not including poorly annotated proteins both decreases the runtime and increases the specificity of SeqScreen functional annotations. SeqScreen then runs Centrifuge, a novel tool for quick and accurate taxonomic classification of large metagenomic datasets. Centrifuge classifications are given higher weights and are always assigned a confidence score of 1.0. SeqScreen always picks the taxonomic rank with the highest score for Centrifuge and assigns it to the sequence. In the case where Centrifuge fails to assign a taxonomic rank to a particular sequence, we assign DIAMOND’s predictions to it. To incorporate DIAMOND’s predictions, we consider all taxonomic ids that are within 1% of the highest bit-score as the taxonomy labels for a sequence (**Supplementary Figure SF1**). The *sensitive* taxonomic classification workflow uses BLASTX and BLASTN for aligning to amino acid and nucleotide databases, respectively. For BLASTX, we again use our reduced version of the UniRef100 database (**Supplementary Data SD1**). BLASTN results are processed through outlier detection to identify which of the top hits are significantly relevant to the query sequence. The *sensitive* mode parameters are set so that if a cut is made, all hits above the cut line are returned; otherwise, all hits are returned. All hits within the outlier detection cutoff (BLASTN) or within 1% (sensitive parameter cutoff=1) of the top bitscore will be saved as the top hits for a given query sequence. Next, all hits reported by BLASTN and BLASTX are sorted by bitscore and listed for a query. Taxonomic IDs are ordered so that BLASTN are reported first, followed by BLASTX. Order-dependent taxonomic assignments will then be based on the first taxonomic ID reported (typically BLASTN hit). Default E-values (--evalue) and max target seqs (--max_target_seqs) for BLASTN and BLASTX are set to 10 and 500, respectively. Since both parameters limit the number of matches to the query sequence, modification of these parameters may be necessary for short and ubiquitous sequences. For BLASTN and BLASTX, the reported confidence values are based on bitscores (bitscore / max bitscore), as inspired by orthology estimation[39].

### SeqScreen workflow #4: Functional Annotation

Using the predicted UniProt IDs and their bit scores from DIAMOND, SeqScreen obtains a list of all predicted UniProt IDs whose bit score is at most 3% less than the highest bit score and compiles all the associated GO terms for each UniProt ID. To assign FunSoCs to each input sequence, we have developed a database which contains a mapping of all UniProt IDs to FunSoCs. The construction of SeqScreen database is described in detail in **Supplementary Data SD1**.

### SeqScreen workflow #5: SeqScreen Reports

Following the computational workflows, SeqScreen produces a tab-separated report file with the predictions of each input sequence as well as an interactive HTML report. The HTML report allows users to search and filter the results based on a variety of criteria such as FunSoC presence, GO term presence, and sequence length. The HTML report is a convenient way to browse the results of large inputs as it loads results in small chunks so that arbitrarily large results can be viewed (**Fig. 3** and **Supplementary Figure SF2**).

**Fig. 3:**
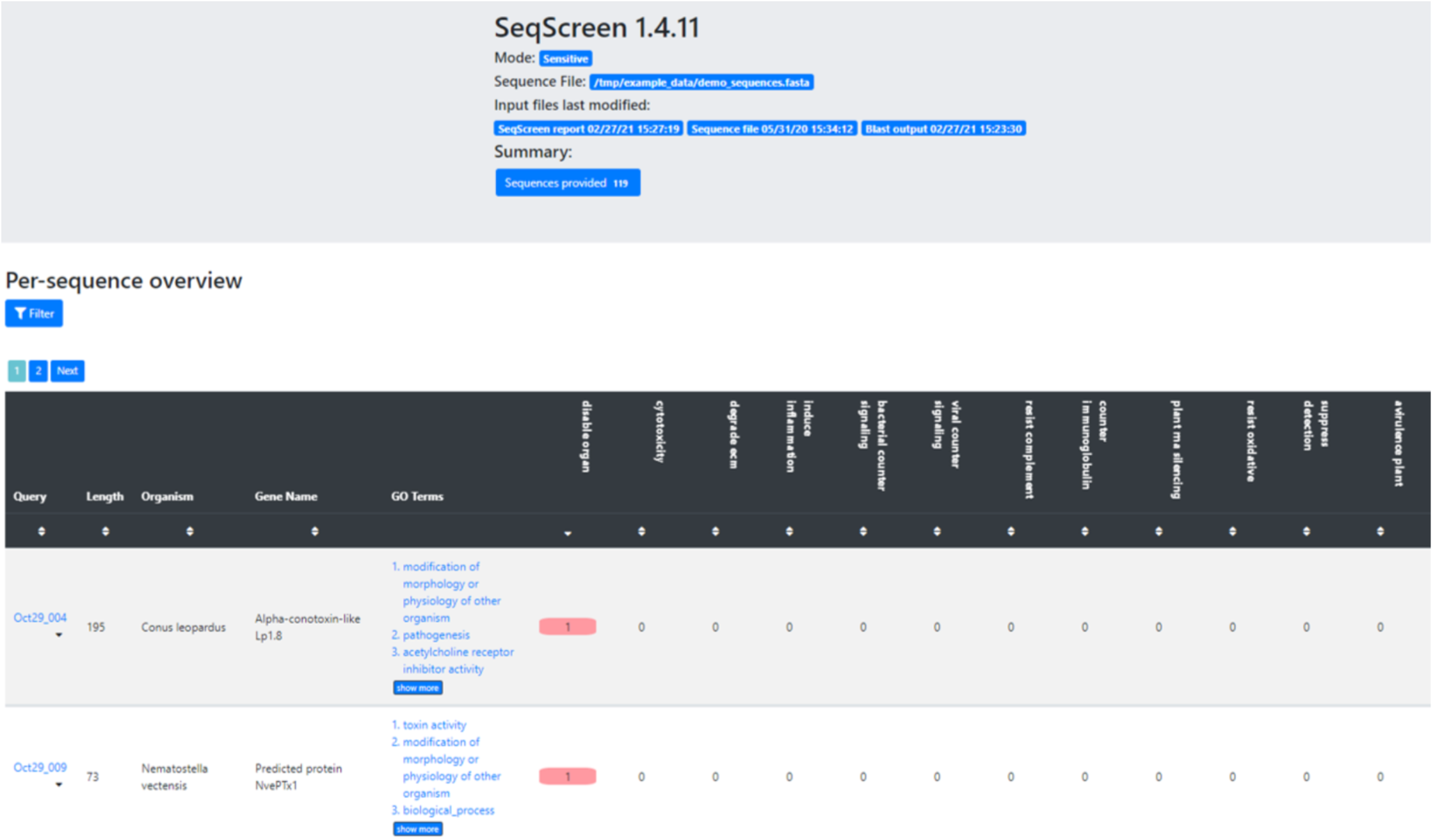
HTML report output from SeqScreen. This is a screenshot of the interactive HTML page that outputs each query sequence in the file, the length, the gene name (if found), and GO terms associated with it. It also outputs the presence (or absence) of each of the 32 FunSoCs by denoting a 1 (or 0) in the given field.

### Ensemble Machine Learning for FunSoC Predictions

FunSoCs encompass sequences involved in the mechanisms of microbial pathogenesis, antibiotic resistance, and eukaryotic toxins (e.g., arachnids, cnidarians, insects, plants, serpents) threatening to humans, livestock, or crops. We identified 32 groups of sequences that could be categorized under the FunSoC framework (**Supplementary Table ST1**) that each protein could potentially be assigned to, thereby indicating pathogenicity. We decided to formulate this as a multi-class, multi-label (i.e., each protein/sequence can be associated with one or more of the 32 FunSoCs) ML classification problem. In order to annotate potentially large numbers of query sequences with FunSoCs, we reasoned that utilizing a lookup table containing pre-predicted FunSoC labels (obtained from the ML models) for the proteins in the UniProt database would enable efficient extraction of labels for corresponding hits from the query to the table. Towards this, we tested 11 ML models (**Supplementary Table ST2**) based on three different strategies that use different feature selection criteria as well as a two-step pipeline that aims to filter proteins that are not associated with any FunSoCs. These models were trained on proteins manually curated and labelled with FunSoCs. For the purposes of our discussion, we show the top three performing models as visualized in **Fig. 4**. To gain a more nuanced understanding of the models’ performances, we considered the average precision and recall of the models on the positive labels specifically, i.e., proteins that were labelled with a “1” (minority class) for a particular FunSoC. This is an important measure to understand how well they learn to predict the minority positive class given the data imbalance which mirrors a practical application of SeqScreen where the expected number of non-pathogenic sequences in a sample is larger than specific pathogenic markers. Our test splits were reflective of this imbalance, for example, the test split for the FunSoC *virulence activity* had 23292 samples labelled “0” and 29 samples labelled “1”. **Table 1** shows the results of different models for each of the metrics. Although the accuracy of the methods is similar, we observed significant differences in the positive label precision and recall. Two Stage Detection + Classification Neural Networks (TS NN) and Two Stage Detection + Classification Balanced Support Vector Classifier (TS Bl.SVC) represented two different ends of the spectrum of precision and recall, the former being more precise (P: 0.88, R: 0.69) and the latter being more sensitive (P: 0.73, R:0.88). We also found that Balanced Support Vector Classifier + Neural Network Classification using Oversampling (Bl. SVC+NN(OS)) represented an intermediate version of the other two models with precision and recall being more balanced (P: 0.87, R: 0.81). The majority vote classifier built on these three classifiers to provide a further improvement in the specificity with a slight loss in terms of recall (P: 0.90, R: 0.82). To get a more detailed perspective of the performance of the models on each of the FunSoCs, we plotted the positive label precision and recall per FunSoC. As seen in **Fig. 5**, the Majority Voting classifier combined the strengths of these individual classifiers to balance precision and recall across these FunSoCs.

**Table 1.** The Accuracy, Exact Match Ratio, Micro and Macro F1 Score, Macro Recall and Precision of the different ML models. The models we considered were Balanced SVC (Feature Selection) + Neural Network Classification using Oversampling (Bl. SVC+NN (OS)), Two Stage Detection + Classification Neural Networks (TS NN), Two Stage Detection + Classification Balanced Support Vector Classifier (TS Bl. SVC), and the Majority Vote Ensemble Classifier (MV ensemble). TS NN had the highest positive label (PL) precision and TS Bl.SVC had the highest positive label (PL) Recall, while Bl. SVC+NN (OS) had the best balance between precision and recall. Majority Vote Ensemble improved on the results of the three classifiers as conveyed by both the high precision and recall the method achieves.

| Model | Accuracy | Exact Match Ratio | Micro F1 Score | Macro F1 Score | Macro Recall | Macro Precision | Mean PL Precision | Mean PL Recall |
| --- | --- | --- | --- | --- | --- | --- | --- | --- |
| Bl. SVC+NN (OS) | 0.9997 | 0.9924 | 0.9859 | 0.8210 | 0.8039 | 0.8716 | 0.8759 | 0.8180 |
| TS NN | 0.9997 | 0.9924 | 0.9359 | 0.6934 | 0.6445 | 0.8011 | 0.8893 | 0.6988 |
| TS Bl.SVC | 0.9996 | 0.9893 | 0.8692 | 0.7047 | 0.8310 | 0.6492 | 0.7382 | 0.8869 |
| MV ensemble | 0.9997 | 0.9934 | 0.9424 | 0.7998 | 0.8016 | 0.8453 | 0.9003 | 0.8273 |

**Fig. 4.**
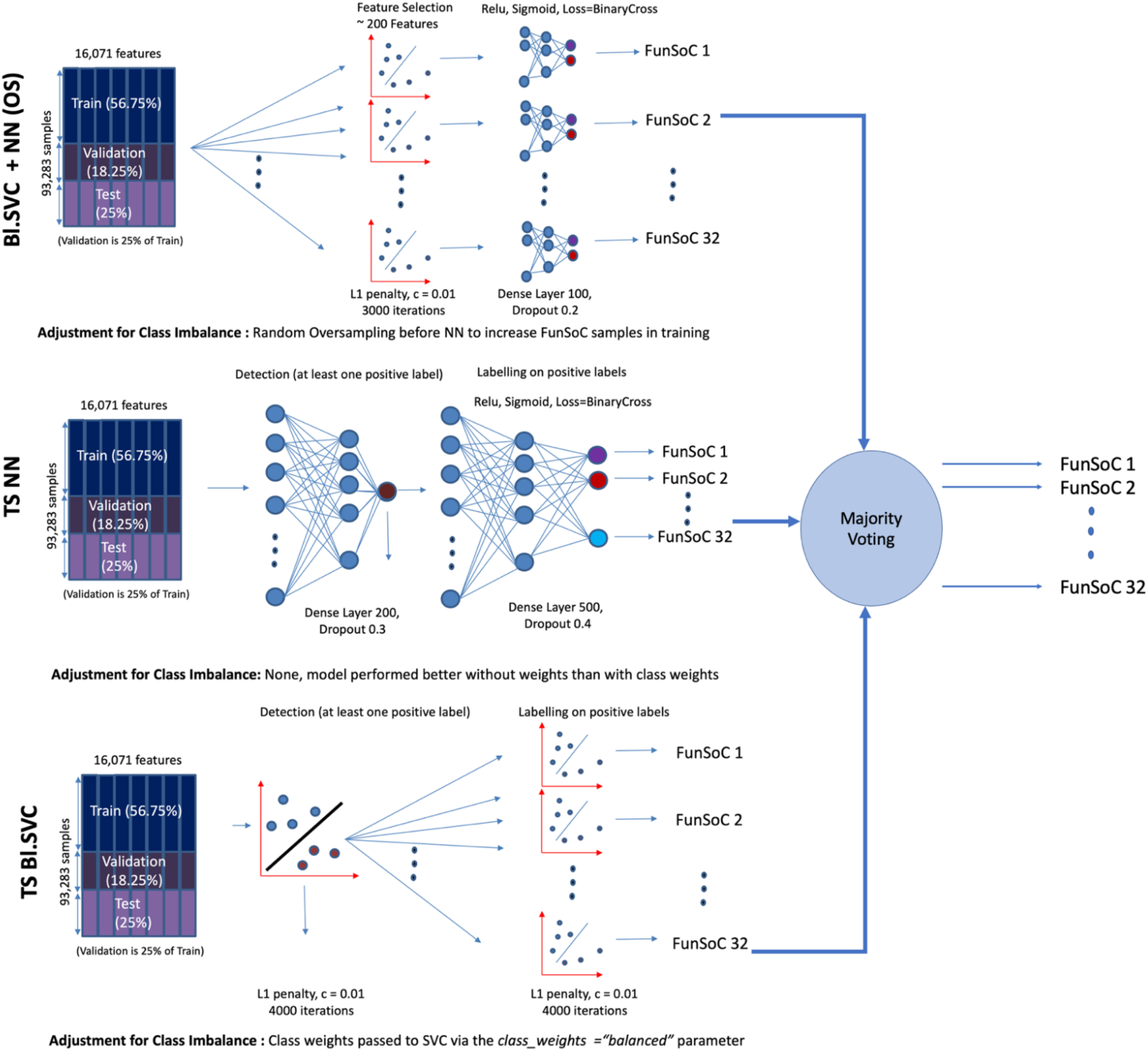
Majority Voting Ensemble Classifier used to create FunSoC Database. The top three models combined are Bl. SVC + NN(OS): Balanced Linear Support Vector Classifier + Neural Networks (Over Sampled), TS NN: Two-stage Neural Network and TS Bl.SVC: Two-stage Balanced Linear Support Vector Classifier. The binary predictions of each of the classifiers over each FunSoC are combined in a majority voting scheme to predict the final labels for the SeqScreen FunSoC database which is then used to annotate query sequences. Training data is split into Train (56.75%), Validation (18.25%) and Test (25%). The two-stage methods fist detect presence of at least one FunSoC and then carry out the multi-class multi-label predictions. Dropouts (Neural Networks) and L1-regularization (Support Vector Classifier) are used to control for overfitting. Two of the models use random oversampling (Bl. SVC + NN(OS), after feature selection) and class weights (TS Bl. SVC) to deal with class imbalance in the training data.

**Fig. 5.**
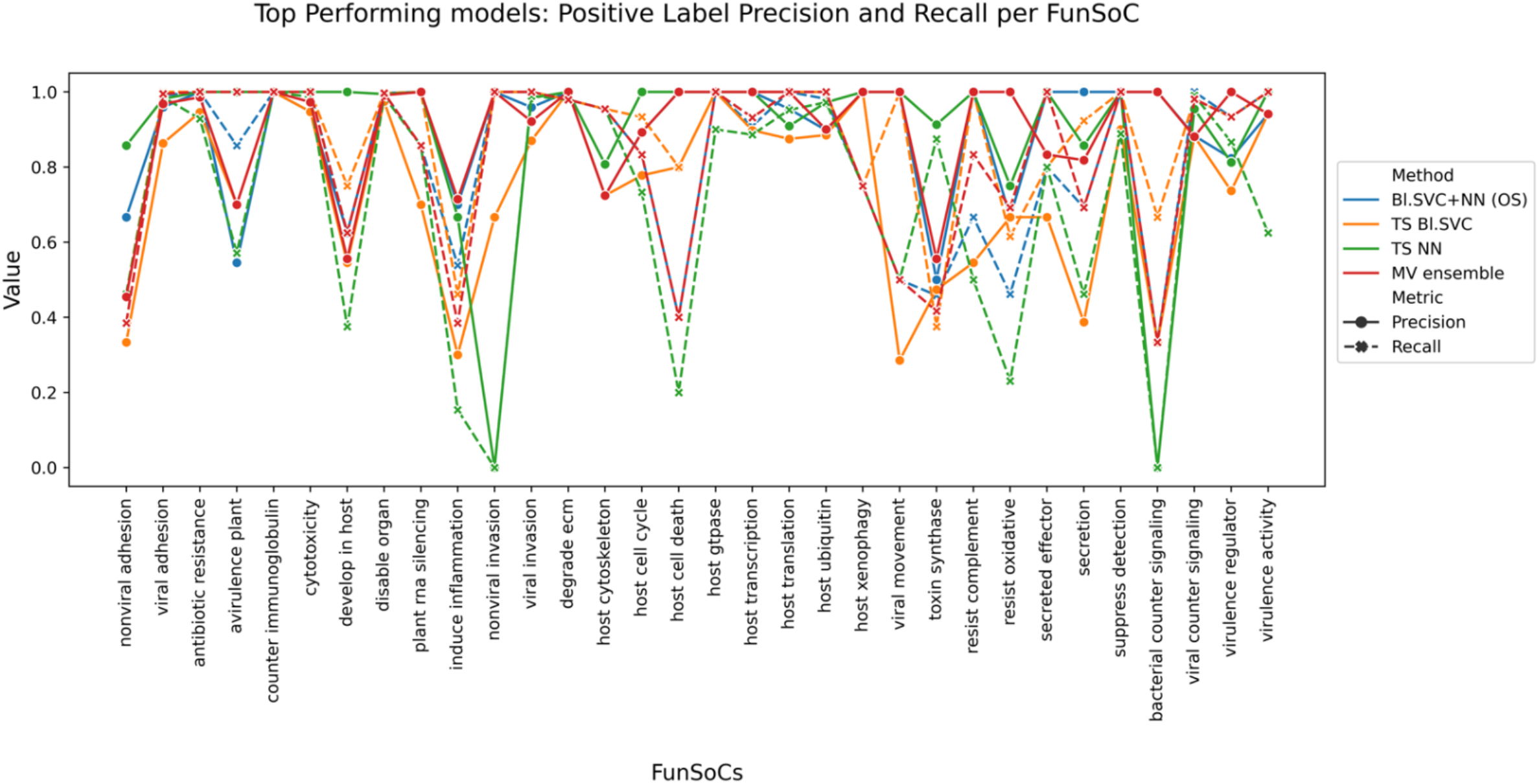
Positive label precision and recall per FunSoc for the four ML models Bl. SVC+NN (OS) (in blue), TS NN (in green), TS Bl. SVC (in orange), and MV ensemble (in red). Precision is in solid lines and Recall is in dotted lines. TS Bl. SVC shows the best overall recall, whereas TS NN consistently has the highest precision across most of the 32 FunSoCs. In hard-to-classify FunSoCs like *nonviral invasion* and *bacterial counter signaling* TS NN performs poorly indicating a model with a high degree of variance. Similarly, TS Bl. SVC suffers from poor precision in most cases. The Majority Vote Classifier improves on the Bl. SVC+NN (OS) and finds an optimal balance between precision and recall across all FunSoCs.

### Use case #1: Screening for known pathogens

We now present a use case with three pairs of hard-to-distinguish bacteria that often confound current metagenomic classification tools to show how SeqScreen analyzes and distinguishes hard-to-classify pathogens. Fig 6. describes the FunSoCs found to be associated with each of the eight bacterial isolate genomes. All isolates showed presence of different antibiotic resistance genes, indicating their ubiquitous presence in most bacteria. In **Fig. 6** (a,b) we show a comparison of the commensal strain of *E. coli K-12 MG1655 versus* the pathogenic strain *E. coli O157:H7*. The two strains showed presence of four FunSoCs, namely *cytotoxicity, secreted effector, secretion*, and *antibiotic resistance*. SeqScreen was able to accurately predict the additional presence of *Shiga toxin subunit B* (*stxB*)[40] in pathogenic *E. coli O157:H7* with the *cytotoxicity* FunSoC and differentiate it from E. *coli K-12 MG1655*. In addition, *E. coli O157:H7* also showed the presence of the *secreted effector protein EspF(U)*, which was labelled with the *secreted effector* and *virulence regulator* FunSoCs. Another example is shown in **Fig. 6** (c,d) where *Clostridium botulinum* and *Clostridium sporogenes* are shown to be differentiated by four specific FunSoCs associated with *C. botulinum*. Though the organisms have a high degree of overall sequence similarity, *C. botulinum* contains the *BotA* toxin which is absent from *C. sporogenes*. We observed the presence of four FunSoCs associated with *C. botulinum*, which included *disable organ, cytotoxicity, degrade ecm* and *secreted effector* associated with hits to the *BotA* and *neurotoxin accessory protein* (*orf-X2)* genes, indicating the presence and the successful detection and annotation of pathogenic genes in *C. botulinum*. In contrast, *C. sporogenes* showed a unique hit to the *secretion* FunSoC, while both organisms were marked with a hit to the *bacterial counter signaling* and *antibiotic resistance* FunSoCs. **Fig. 6** (e,f) shows that FunSoCs can also be used to differentiate between *Streptococcus pyogenes* (Group A Streptococcus, causative agent of Strep throat) and *Streptococcus dysgalactiae* (Group C/G Streptococcus), a near neighbor with pathogenic potential. *S. pyogenes* had the *streptopain* (*speB*) and *exotoxin type H* (*speH*) genes associated with the *induce inflammation* FunSoC, whereas *S. dysgalactiae* had the *immunoglobulin G-binding protein* (*spg*) gene with the *counter immunoglobulin* FunSoC, thereby differentiating it from *S. pyogenes*. Both bacteria showed presence of *cytotoxicity, secretion*, and *antibiotic resistance*. In addition to pathogens, we show in **Fig. 6 (g,h)** that the FunSoC based framework can also capture well-characterized commensals like *Streptococcus salivarius* and *Lactobacillus gasseri*. We see that both these bacteria reported the least number of FunSoCs, validating the negative control experiment. *S*.*salivarius* contained a hit the *secretion* FunSoC from genes encoding competence proteins. In differentiating near neighbor pathogens, SeqScreen selectively annotated regions in genomes that contributed to pathogenicity across various categories.

**Fig. 6.**
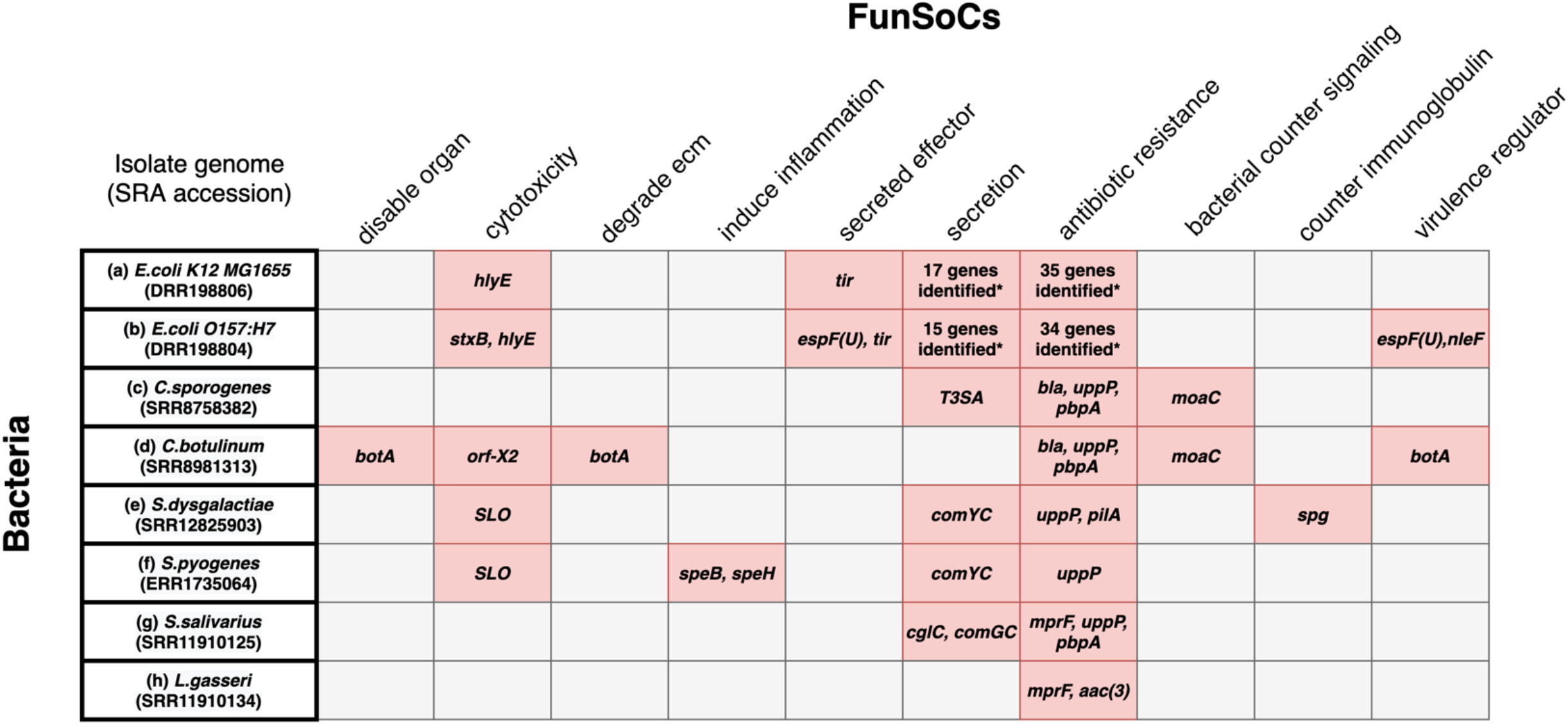
Pathogen identification of hard-to-classify pathogens: FunSoCs Assigned to Genes by SeqScreen. Abbreviated gene names are listed in pink cells if at least one read from the gene had a UniProt e-value < 0.0001, was assigned a FunSoC, and was from the expected genus (i.e., Escherichia or Shigella, Clostridium, Streptococcus, Lactobacillus). FunSoCs with at least one gene that met the criteria for detection in at least one isolate were included in the table. The removal of genes from genera that were not expected in these bacterial isolates allowed for removal of genes that were likely derived from likely contaminating organisms (e.g., PhiX Illumina sequencing control). An expanded table for cells denoted by (*) and complete gene names are listed within each cell in **Supplementary Table ST3**. (a and b) *E. coli O157:H7* is shown to have presence of the *shiga toxin (stxB)* as seen in the *cytotoxicity* FunSoC, as well as an additional hit to the *secreted effector protein* (*espF(U*)), labelled with *secreted effector* and *virulence regulator* FunSoCs, compared to *E*.*coli K12 MG1655*. (c and d) *C. botulinum* showed four distinct FunSoCs (*disable organ, cytotoxicity, degrade ecm* and *virulence regulator*) and presence of the *botA* and *orf-X2* genes compared to *C. sporogenes*. (e and f) *S. pyogenes* showed presence of the *induce inflammation* FunSoC in contrast to the near neighbor pathogen *S. dysgalactiae* with the *counter immunoglobulin* FunSoC. (g and h). *S. salivarius* and *L. gasseri* are well-known commensals that are generally considered harmless. Both show presence of antibiotic resistance genes, while *S. salivarius* also contains some genes associated with *secretion*. The commensals have hits to the least number of FunSoCs.

In addition to FunSoCs assignments, we evaluated how existing alignment approaches handle the identification of pathogen near neighbors. To motivate our experiments, we initially considered the widely used BSAT list to triage isolates (see Methods section on SeqMapper), as it is representative of a current strategy for pathogen screening approaches in the DNA synthesis industry. We mapped *C. sporogenes* (SRR8758382) reads against the BSAT database using the Bowtie2 module of the SeqMapper workflow, and 98.28% of the reads hit to *C. botulinum*. The high percentage of hits to *C. botulinum* underlines the shortcoming of simplistic triaging methods to accurately differentiate between near neighbors and pathogens. We further considered popular taxonomic classifiers to analyze how accurately near neighbor pathogens were separated. We compared the results of six different tools, Mash dist[41], Sourmash[42], PathoScope[8], Kraken2[43], MetaPhlAn3[44], KrakenUniq[45] and Kaiju[46], with the following three pairs of near-neighbors and pathogens: *E. coli K-12 MG1655 and E. coli O157:H7, C. sporogenes and C. botulinum*, and *S. dysgalactiae and S. pyogenes*. **Table 2** shows the results of running the taxonomic tool on these bacteria with their complete databases and the top hits for each are reported. Strain level differences between the two *E. coli* near neighbor was hard for almost all the tools to distinguish. Kaiju and MetaPhlAn3 could only predict *E. coli* at species level for both strains, and since those tools were designed to only report down to the species level, strain-level pathogenicity will always be missed. Kraken2 incorrectly predicted non-pathogenic *E. coli K-12 MG1655* as the pathogenic strain *E. coli O157:H7*. PathoScope and KrakenUniq incorrectly predicted the non-pathogenic *E. coli K-12 MG1655* strain as *E. coli BW2952* and *E. coli O145:H28*. Mash dist and Sourmash were the only tools that reported the true *E. coli K-12* strain. The tools performed considerably better when predicting for *E. coli O157:H7*, as Mash dist, Sourmash, PathoScope, Kraken2 and KrakenUniq were able to predict the strain correctly. When considering the two *Clostridium* near neighbors, PathoScope, Kraken2 and KrakenUniq misclassified *C. sporogenes* as *C. botulinum*. In contrast, *C. botulinum* was incorrectly called *C. sporogenes* by Mash dist, Sourmash and MetaPhlAn3. While predicting for the *Streptococcus* near neighbors, all tools predicted *S*.*pyogenes* correctly and only PathoScope misclassified *S. dysgalctiae* as *S. pyogenes*, while other tools called it accurately. In summary, our experiments demonstrated that none of the tools were able to correctly predict all pathogens and near neighbors at the species and strain levels. SeqScreen provides a more detailed framework beyond species or strain-level taxonomic classifications to aid the user in interpreting the pathogenicity potential of a query sequence, including exact protein hits, GO terms, multiple likely taxonomic labels with confidence scores, and FunSoC assignments.

**Table 2.**
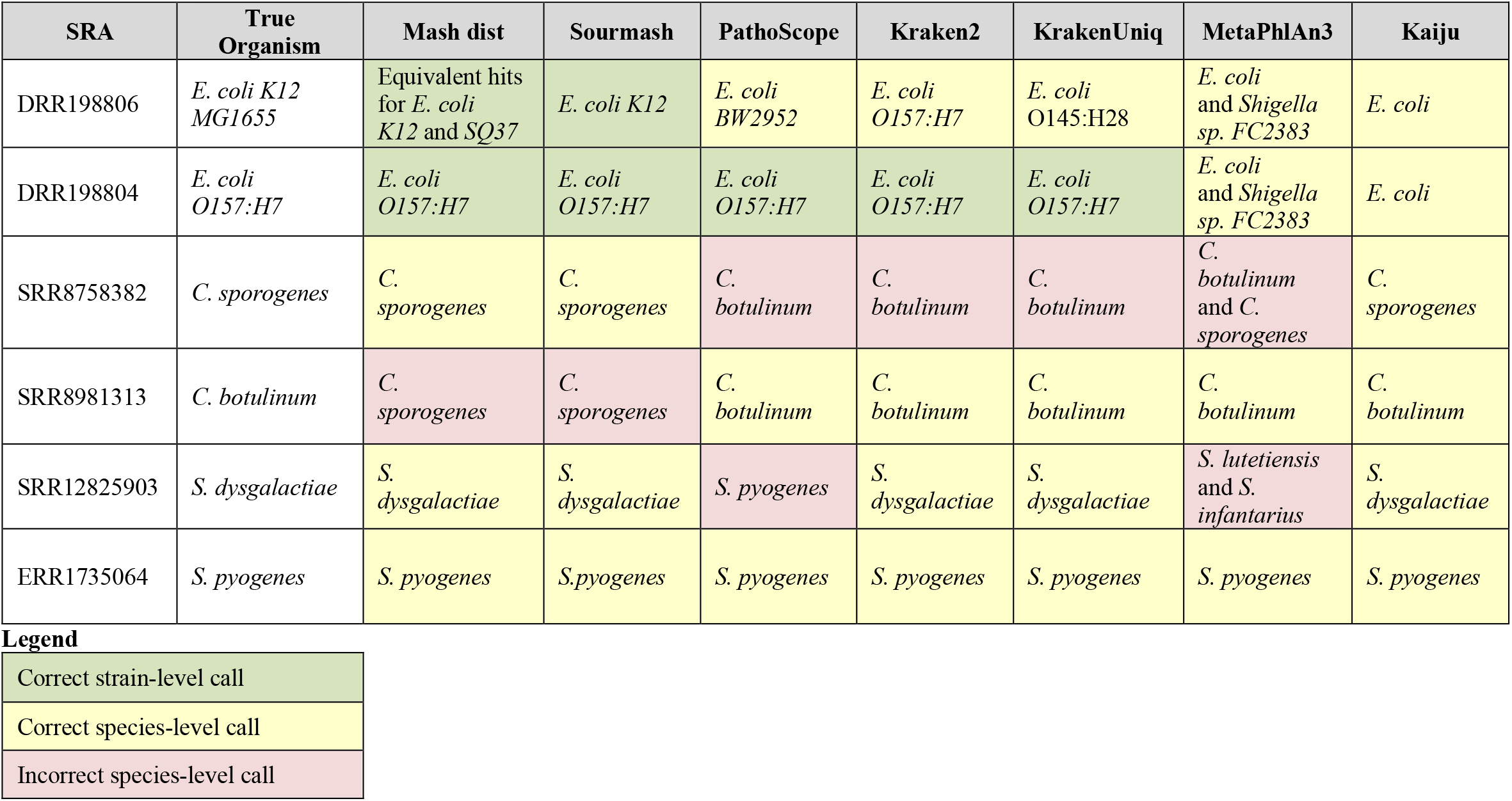
Pathogen and near neighbor classification. **SRA** represents the SRA id of the sample, **True Organism** represents the actual bacterial strain or species, and the remaining columns indicate the results for the indicated method using the parameters detailed in the Methods section. Green cells indicate that the tool assigned a correct strain-level call, yellow indicates a correct species-level call, and red indicates an incorrect species-level call. The following tools and databases were run: Mash dist (RefSeq 10k), Sourmash (RefSeq + GenBank), PathoScope (PathoScope DB), Kraken 2 (Mini and full Kraken2 DB produced the same results), KrakenUniq (MiniKraken 8GB), MetaPhlAn3 (default) and Kaiju (index of NCBI nr + euk). The *E. coli* strains were challenging for most tools. The pathogenic *E. coli O157:H7* was correctly called by Mash dist, Sourmash, PathoScope, Kraken2 and KrakenUniq. MetaPhlAn and Kaiju could only make a species level assignment. In contrast, the commensal *E. coli K12 MG1655* was the most challenging as only Mash dist and Sourmash got the strain level assignment correct. MetaPhlAn3 and Kaiju could make only species level assignments, and PathoScope, Kraken2, and KrakenUniq called it as strains *E. coli BW2952, E. coli O157:H7*, and *E. coli O145:H28*, respectively.Even with a complete database, *C. sporogenes* was wrongly classified as *C. botulinum* by PathoScope, Kraken2, and KrakenUniq. Mash dist, Sourmash, and Kaiju predicted *C. sporogenes* correctly while MetaPhlAn3 was ambigous. *C. botulinum* was incorrectly classified as *C. sporogenes* by Mash dist, Sourmash, *S. dysgalactiae* was predicted as *S*.*pyogenes* by PathoScope. All tools correctly called *S. pyogenes*.

| SRA | True Organism | Mash dist | Sourmash | PathoScope | Kraken2 | KrakenUniq | MetaPhlAn3 | Kaiju |
| --- | --- | --- | --- | --- | --- | --- | --- | --- |
| DRR198806 | *E. coli* K12 MG1655 | Equivalent hits for *E. coli* K12 and SQ37 | *E. coli* K12 | *E. coli* BW2952 | *E. coli* O157:H7 | *E. coli* O145:H28 | *E. coli* and *Shigella* sp. FC2383 | *E. coli* |
| DRR198804 | *E. coli* O157:H7 | *E. coli* O157:H7 | *E. coli* O157:H7 | *E. coli* O157:H7 | *E. coli* O157:H7 | *E. coli* O157:H7 | *E. coli* and *Shigella* sp. FC2383 | *E. coli* |
| SRR8758382 | *C. sporogenes* | *C. sporogenes* | *C. sporogenes* | *C. botulinum* | *C. botulinum* | *C. botulinum* | *C. botulinum* and *C. sporogenes* | *C. sporogenes* |
| SRR8981313 | *C. botulinum* | *C. sporogenes* | *C. sporogenes* | *C. botulinum* | *C. botulinum* | *C. botulinum* | *C. botulinum* | *C. botulinum* |
| SRR12825903 | *S. dysgalactiae* | *S. dysgalactiae* | *S. dysgalactiae* | *S. pyogenes* | *S. dysgalactiae* | *S. dysgalactiae* | *S. lutetiensis* and *S. infantarius* | *S. dysgalactiae* |
| ERR1735064 | *S. pyogenes* | *S. pyogenes* | *S. pyogenes* | *S. pyogenes* | *S. pyogenes* | *S. pyogenes* | *S. pyogenes* | *S. pyogenes* |

Legend
|  |
| --- |
| Correct strain-level call |
| Correct species-level call |
| Incorrect species-level call |

### Use case #2: Screening for novel pathogens

To highlight the advantage of using SeqScreen’s FunSoC based pathogen detection pipeline in contrast to relying on taxonomic labels, our next set of experiments and results evaluated how the absence of the exact set of species or strain entries in the database corresponding to the bacterial genome query would impact the classifications by these tools. This was done to simulate a query of a novel pathogen genome by removing the entries corresponding to the query bacterial genome from the database. We chose two tools for this experiment, Mash dist and PathoScope, as modifying their databases for this experiment was readily achievable and both performed well in the previous use case. **Table 3** shows the results of the classifiers using these modified databases. As expected, the closest near neighbor of the query genome is selected when a pathogen is not present in the database, which while representing the expected behavior, is not suitable for sensitive flagging of pathogenic sequences. As is the case with the complete databases, both the tools misclassified the *E. coli* strains with PathoScope only being able to classify *E. coli K-12 MG1655* at species level and Mash dist instead reporting a hit to the pathogenic *E. coli O16:H48* strain. For the *Clostridium* species, both the tools called the pathogen as its non-pathogenic near neighbor, emphasizing the difficulty of identifying these pathogens in a simulated novel pathogen environment. In the case of *Streptococcus, S. dysgalactiae* was classified as *S. sp. NCTC 11567* by Mash dist and *S. intermedius* by PathoScope, whereas *S. pyogenes* was classified as its near neighbor *S. dysgalactiae* by Mash dist and *S. infantarius* by PathoScope. In contrast, as seen in Fig. 4, retaining genus specific hits from SeqScreen was sufficient to observe functional differences between the near neighbor pathogens. This experiment showed that current approaches may still fail to separate near neighbor pathogens and hence a novel FunSoC-based functional framework could help fill the gap and capture sequence level pathogenic markers.

**Table 3.**
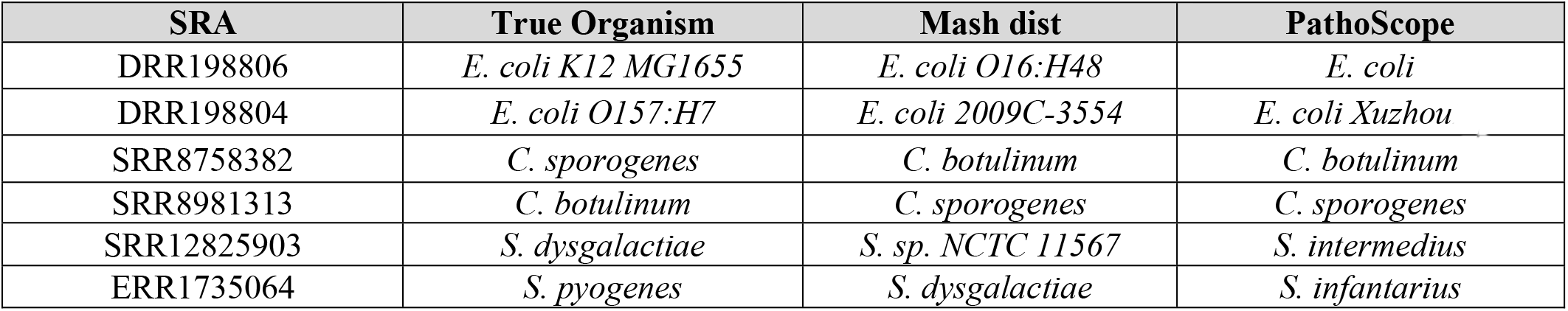
Simulating a novel pathogen. Mash dist and PathoScope were run on pathogen sequences and their near neighbors with the corresponding truth species removed in their respective databases to simulate an example of classifying a novel pathogen not in the database. **SRA** represents the SRA id of the sample, **True Organism** represents the actual bacterial strain or species, **Mash dist** represents the Mash results on each of the samples (with the truth organism species or strain removed from its sketch database), and Pathoscope represents the PathoScope results on each of the samples (with the truth organism species or strain removed from its database). In three of the cases, *C. sporogenes, C. botulinum* and *S. pyogenes*, Mash dist classified the organism as it near neighbor - *C. botulinum, C. sporogenes and S. dysgalactiae*, respectively. *S. dysgalactiae* was classified as *S. sp. NCTC 11567* whereas the commensal *E. coli K12* and pathogenic *E. coli 0157:H7* were classified as *E. coli O16:H48* and *E. coli 2009C-3554*, respectively. PathoScope only classified two pathogens, *C. sporogenes* and *C. botuinum*, as their nearest neighbor counterparts. *S. dysgalactiae* was classified as *S. intermedius*, whereas *S. pyogenes* was classified as *S. infantarius. E. coli K12* was only classified at the species level, while the pathogenic strain *E. coli O157:H7* was classified as *E. coli xuzhou21*.

| SRA | True Organism | Mash dist | PathoScope |
| --- | --- | --- | --- |
| DRR198806 | *E. coli* K12 MG1655 | *E. coli* O16:H48 | *E. coli* |
| DRR198804 | *E. coli* O157:H7 | *E. coli* 2009C-3554 | *E. coli* Xuzhou21 |
| SRR8758382 | *C. sporogenes* | *C. botulinum* | *C. botulinum* |
| SRR8981313 | *C. botulinum* | *C. sporogenes* | *C. sporogenes* |
| SRR12825903 | *S. dysgalactiae* | *S. sp. NCTC 11567* | *S. intermedius* |
| ERR1735064 | *S. pyogenes* | *S. dysgalactiae* | *S. infantarius* |

### Use case #3: Screening human clinical samples for an unknown pathogenic virus

As a final use case to further illustrate SeqScreen’s ability to identify pathogenic sequences in clinical samples, we ran SeqScreen on the sequencing data obtained from the peripheral blood mononuclear cells (PBMC) of three COVID-19 patients and three healthy patients as reported in the study by Xiong et al[47]. We reasoned that the samples from COVID-19 patients should contain certain reads with functional markers that would indicate presence of the SARS-CoV-2 virus. To better understand SeqScreen’s application in analyzing clinical samples for unknown pathogenic viruses, we chose to run an older version of SeqScreen (v1.2) on these samples, retaining the same analysis functionality with a database that predated the COVID-19 pandemic and the inclusion of SARS-CoV-2 virus. This was done for two main reasons. First, we wanted to evaluate SeqScreen’s ability to retrieve functional pathogenic information by simulating an experiment with an unknown virus along with a database that did not contain the causative virus. Second, we wanted to highlight SeqScreen’s ability to detect GO terms and FunSoCs directly from metatranscriptomes of clinical samples with low levels of the novel pathogen. For this study, we focused on GO terms that were specific to the COVID-19 samples and viral proteins (i.e., GO terms that were not assigned to bacterial, eukaryotic, or archaeal proteins or observed in the healthy controls). Only three GO terms met these criteria within one of the COVID-19 samples (CRR119891). All three of the GO terms, suppression by virus of host ISG15 activity (GO:0039579), induction by virus of catabolism of host mRNA (GO:0039595), and suppression by virus of host NF-kappa B transcription factor activity (GO:0039644) were indicative of SARS-CoV-2 virus activity. SeqScreen assigned replicase polyprotein 1ab from Bat coronavirus 279/2005 (UniProt ID: P0C6V9, e-value: 5.8e-29) to one sequence read and reported these three GO terms in sample CRR119891. Searching for other coronavirus taxonomic assignments in that sample revealed one additional read that SeqScreen assigned to spike glycoprotein from Bat coronavirus HKU3 (UniProtID: Q3LZX1, e-value: 1.3e-09). No other coronavirus reads were identified in the samples, consistent with the report from the original publication in Xiong et al that very few to no SARS-CoV-2 reads were identified in the PBMC samples. In the SeqScreen v1.2 database, the associated FunSoC with the replicase polyprotein 1ab was evasion and the FunSoCs predicted for the spike protein were adhesion and invasion, which reflect the biological functions of the two proteins and indicate presence of virulence. To compare SeqScreen v1.2 results to another tool, we ran HUMAnN2[48] on the six PBMC metatranscriptomes to check for presence of virulence markers and pathways. The HUMAnN2 results did not point to any evidence for presence of COVID-19 specific markers in this sample nor the others (**Supplementary Data SD2**), which is expected given the focus of the tool on reporting enriched genes and pathways, rather than rare pathogenic sequences. As SeqScreen extensively characterizes individual short protein-coding sequences and is geared towards identifying functional markers of pathogenicity, it can sensitively detect trace amounts of pathogenic signal in clinical samples. The reads identified as SARS-CoV-2 were confirmed to such when aligned to the database containing SARS-CoV-2 using BLAST[28] as seen in **Fig. 7**.

**Fig. 7:**
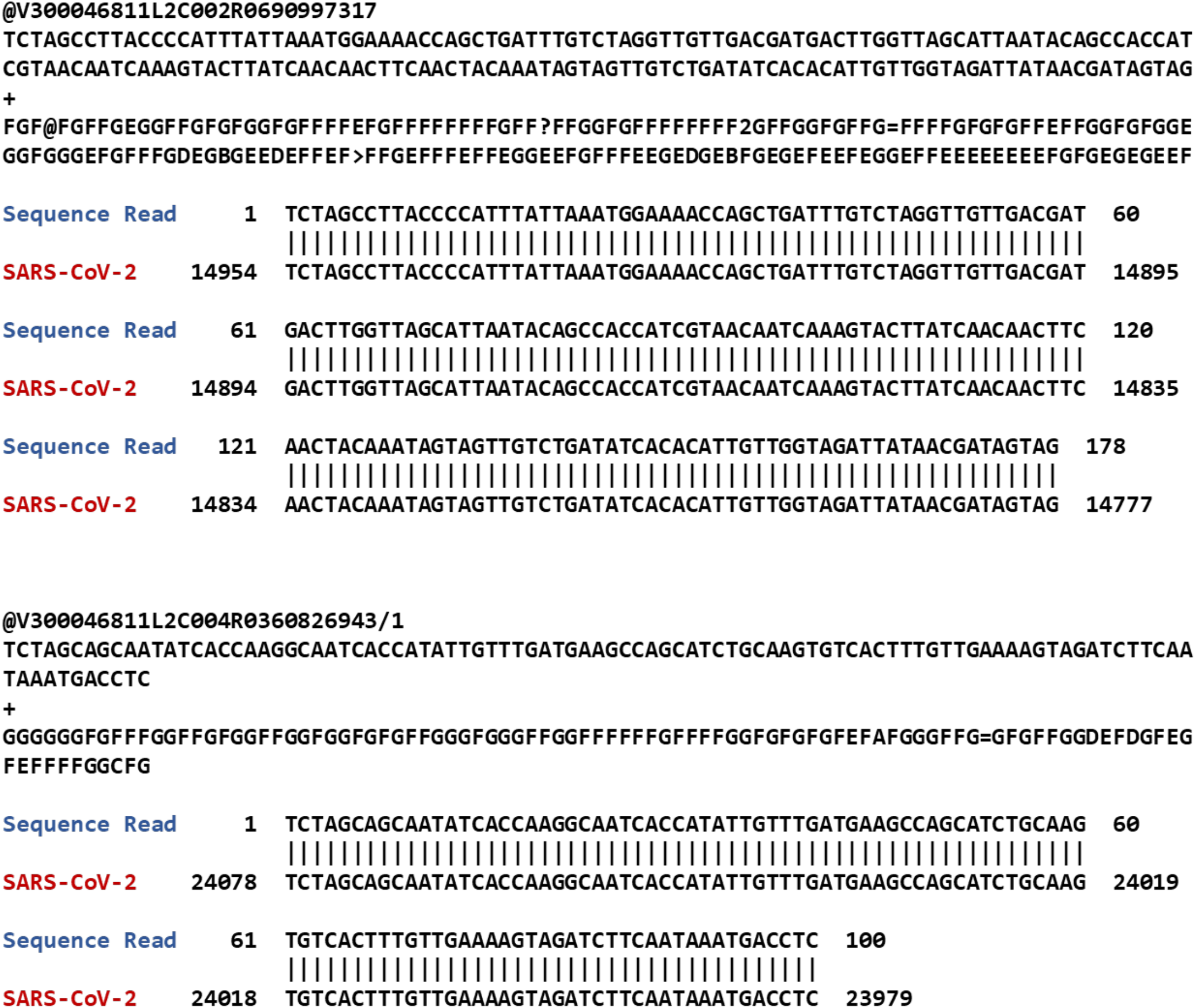
Two trimmed fastq reads identified by SeqScreen v1.2 in sample CRR119891, and their alignment to the SARS-CoV-2 genome: BLAST results of the specific reads in the samples of COVID-19 infected patients from Xiong et al.[47] against SARS-CoV-2 genome. These reads were identified by SeqScreen as belonging to SARS-CoV-2 (without it being present in the SeqScreen DB).

## Discussion

The challenge of pathogen identification and detection from sequence level features is significant and requires a nuanced, multi-layered approach. A given genus or species often includes both pathogenic and non-pathogenic strains. These may not be well-defined by taxonomic considerations[4] since sequences with similar taxonomic labels may contain pathogenic elements as well as non-pathogenic markers. Even at the strain level, addition or subtraction of a single gene may affect the overall pathogenicity of the microbe. SeqScreen provides a novel approach to this important problem and focuses on read-level analyses that facilitate the detection of low abundance pathogenic markers from metagenomic samples. Not only does SeqScreen analyze partial and full-length genes specific to FunSoCs, sequences annotated with a subset of high-confidence FunSoCs can be analyzed to detect pathogenic presence in the sample. Taxonomic classifiers often are ambiguous about similar pathogens and near neighbors within the same genus or species, such as commensal *E. coli K-12 MG1655* and pathogenic *E. coli O157:H7*, as well as *C. botulinum* and *C. sporogenes*, and *S. dysgalactiae* and *S. pyogenes*. We show that FunSoCs can be used as unique signatures to distinguish these pairs. We also saw that commensal bacteria such as *L. gasseri* had no FunSoCs associated with it, other than antibiotic resistance which has been previously reported[49], validating our negative control and highlighting SeqScreen’s ability to accurately identify commensals. Note, several the commensals analyzed in this study contain genes that can cause infection in humans, but these microbes are rarely disease-causing agents.

A notable, novel feature of SeqScreen for pathogen detection and characterization is the addition of FunSoCs as a labeling system for each sequence in the query. FunSoCs are molecular activities of pathogens that contribute to its pathogenesis in human, crop, or livestock hosts. Using controlled vocabularies and other data mined from popular protein databases, we showed that our models can capture FunSoCs with a high level of precision. To improve the balance between precision and recall over most of the FunSoCs, we proposed a majority voting ensemble classifier. SeqScreen utilizes a lookup table created by classifying all UniProt proteins using the ensemble classifier to annotate query sequences with FunSoCs.

SeqScreen’s FunSoC curations are not the first attempt to collate sequences of concern in a specific computational framework and/or database. Prior efforts such as the Virulence Factor Database (VFDB), Pathosystems Resource Integration Center (PATRIC)[50], and Pathogen-Host Interaction database (PHI-base)[51] all offer resources for identification of virulence factors and pathogenic sequences. VFDB is a database of virulence factors that have been widely used but is limited due to many of the sequences not having clearly available annotations or justification for their pathogenic status. PATRIC primarily focused on annotation of isolates/pathogens but not individual sequences, and PHI-base describes the pathogen-host interactions but does not focus on pathogenic effects on the host. SeqScreen was designed to specifically overcome some of the major limitations through an iterative ensemble learning framework that leverages functional information combined with curations to identify FunSoCs

Our experimental results underscore the importance of using a function-based framework in contrast to the prevailing taxonomy-based classifiers and pathogen detection tools. SeqScreen’s FunSoC based pathogen detection approach is sensitive to specific gene-based differences between closely related strains and accurately identifies pathogenic markers. Out of the tools we evaluated, only Kaiju was able to accurately distinguish all the near neighbors from pathogens at the species level. The protein-based classification strategy used by Kaiju is different from other k-mer based tools, but similar to SeqScreen’s functional based characterization framework, indicating the advantages of using the functional units in proteins to identify pathogens. SeqScreen provides an advantage in that it also reports the most likely strain-level assignments and protein-specific functional information for each sequence, including GO terms and FunSoCs, to accurately identify pathogenic markers in each sequence without relying solely on taxonomic markers. We also observed through inspecting the FunSoC lookup table that SeqScreen preserves FunSoC labels even when the proteins are distantly related (up to 40% sequence similarity). Hence, the FunSoC abstraction represents a robust framework to detecting novel pathogens as it does not rely on specific taxonomic labels in the database but on learning latent features that connect similar pathogenic makers. SeqScreen also provides a more detailed framework beyond species or strain-level taxonomic classifications to aid the user in interpreting the pathogenicity potential of a query sequence, including exact protein hits, GO terms, multiple likely taxonomic labels with confidence scores, and FunSoC assignments.

The task of mapping biological (e.g., functional annotations) and textual features (e.g., keywords and abstract metadata) to these FunSoCs is non-trivial for three reasons. The first concerns identifying from the literature a sufficiently large training set of sequences associated with each FunSoC. Second, variability in annotation across subject matter experts and inconsistencies in database annotations often makes it challenging to incorporate relevant features. Third, the amount of labelled data available per FunSoC is disproportionate which makes accurate multi-label and multi-class classification difficult. Also, the positive labels are far fewer when compared to the negative labels making the accurate prediction of positive labels non-trivial due to class imbalance.

One known limitation of SeqScreen is that it heavily depends on annotated sequences for identification of FunSoCs. As of April 2021 (UniProt release 2021_02), there are 1.5 million proteins with evidence at the protein or transcript level (less than 0.75%), with 64 million proteins with functions inferred from homology and over 212 million proteins total. Through several years of curating, our team was able to characterize thousands of proteins specific to pathogenic function, augmenting information contained in UniProt, and enabling robust pathogenic sequence screening of sequences of high concern. However, coordinated community efforts are needed to further extend out and improve annotation quality of proteins in these key databases. We also note that while we have shown SeqScreen to be an accurate pathogen detection tool, explicitly identifying and labelling pathogens is not possible with only FunSoC information, as seen in Fig. 4, and the presence of genes underlying the FunSoC annotations should be considered when interpreting results. SeqScreen identifies and flags sequences having functions of concern (or FunSoCs) but stops short of performing pathogen identification, as it was designed to only characterize individual DNA sequences. In future work, we aim to extend our FunSoC-based machine learning (ML) framework towards pathogen identification by analyzing sequences at the whole genome level.

Finally, while SeqScreen can accurately screen oligonucleotides and short DNA sequences for FunSoCs, large metagenome-scale pathogen analysis is still an open challenge. Currently, the accuracy and sensitivity of SeqScreen annotation comes at a substantial cost of runtime and memory requirements compared to other tools and pipelines. To address this, one possible solution is to use a read or database subsampling method such as RACE[52] that may be able to preserve the full complement of taxonomic and functional diversity while drastically reducing runtime.

## Conclusions

SeqScreen describes a novel, comprehensive sequence characterization and pathogen detection framework based on a multimodal approach that combines conventional alignment-based tools, machine learning and expert biocuration to produce a new paradigm for novel pathogen detection tools beneficial to both synthetic DNA manufacturers and microbiome scientists alike. SeqScreen is the first open-source, modular framework for transparent and collaborative research to improve DNA screening practices beyond simple screens against BSAT agents and toxins.

## Methods

### Pipeline Implementation

The SeqScreen pipeline is implemented as a modular architecture combining various individual workflows for taxonomic and functional characterization as well as identification of Functions of Sequences of Concern (FunSoCs) in short DNA sequences. The pipeline is implemented using Nextflow for scalable and reproducible deployment and the scripts are written in Perl and Python. The five main workflows available as part of SeqScreen are (i) Initialization (fasta verification) (ii) SeqMapper (Identification of BSAT agents) (iii) Protein and Taxonomic identification (iv) Functional annotation (v) FunSoC identification and SeqScreen report generation. Further information on databases, dependencies and parameters can be found at GitLab: https://gitlab.com/treangenlab/seqscreen/-/wikis/home. The modules used depend on the mode (default or sensitive) that SeqScreen is run. In the slower sensitive mode, BLAST(N/X) approaches are used to get an accurate protein and taxonomic identification and functional annotations. In contrast, default mode is faster as it uses DIAMOND (--evalue 10 –block-size 200 –more-sensitive) for protein identification. The taxonomic classification workflow in this mode combines both centrifuge and DIAMOND results. In addition to different modes, SeqScreen also has optional modules like HMMER which can be activated with a flag (--hmmscan) which runs the sequence against the Pfam HMMs. To increase the efficiency of analysis, SeqScreen also supports multithreading as well as SLURM execution (--slurm) for runs on High Performance Computing (HPC) nodes. FunSoCs are assigned to sequences by transferring labels from protein hits. The output includes a report in TSV format that captures the taxonomic and functional as well as FunSoC annotations for each read in the sample. SeqScreen also provides a HTML view of the FunSoCs for each of the sequences in the sample with additional filters for users to view and select sequences and/or FunSoCs of interest.

### Functional Benchmarking

Data for the functional benchmarking was downloaded from the CAFA website (https://www.biofunctionprediction.org/cafa/). The CAFA 3[53] training data was downloaded from the website (https://www.biofunctionprediction.org/cafa-targets/CAFA3_training_data.tgz). From the training set, a subset of 250 proteins having appropriate lengths (at least 200 aa) were chosen for the benchmarking. A set of (250) proteins of sub-lengths 34 aa, 50 aa, 67 aa and 80 aa was derived from this set of proteins for sub-lengths benchmarking. To create the sub-lengths for the respective proteins, we randomly selected a starting residue from each of the 250 proteins and considered the stretch of residues up to the desired lengths as the sub-protein. The proteins were then run through each of the tools: PANNZER2[54], eggNOG-mapper[55] and DeepGOPlus[56]. Further details about the dataset, tools and commands and databases the tools were run with are shown in the **Supplementary Data SD3.1** and **Supplementary Table ST4**.

### Taxonomic Benchmarking

Seven simulated datasets used in previous tool benchmarking and comparison studies were considered for benchmarking[57,58]. These reflected characterized real metagenomes found in various environments like human (e.g., buccal, gut) and in the natural or built environment (e.g., city parks/medians, houses, soil, subway), using the same methodology. All reads were 100-bp (Illumina) and simulated using ART [59] at 30X coverage and post-processed to remove ambiguously mapped reads at the species levels using MEGAN[60]. The reads thus obtained map unambiguously to a single species in the RefSeq database. SeqScreen’s performance on taxonomy and additional information can be found in **Supplementary Data SD3.2** and **Supplementary Table ST4**.

### Ensemble Machine Learning for FunSoC Prediction

One of the major applications of SeqScreen is its ability to combine functional and taxonomic information for pathogen detection. To assign pathogenic functions to query sequences in each sample, SeqScreen labels relevant sequences with FunSoCs. Each FunSoC captures a process contributing either to pathogenesis or countermeasure resistance. Proteins representing the FunSoCs were identified primarily through literature review with some database perusal (VFDB, PHI-base). The expert human biocurators developed queries using terms from controlled vocabularies and in specified UniProt fields to obtain sequence sets for each FunSoC. Examples of these UniProt queries are provided in **Supplementary Data SD4**. After initial formulation with UniProt queries, the biocurator FunSoC annotations were verified through manual literature reviews thereby maximizing the number of sequences specific to the FunSoC category while eliminating false positives. An updated database of SeqScreen biocurated FunSoCs is maintained in **Supplementary Data SD5**. The proteins of each FunSoC were then used as a training set. The training set sizes for each FunSoC ranged from 4,722 for *disable organ* to 24 for *counter immunoglobulin*. These also included proteins that had annotation scores less than 3, which were pruned out in the preprocessing step to get high-quality labelled training data. We used these proteins as the training dataset for our Machine Learning models to capture underlying mappings between the sequence features and FunSoCs. Each of the curated proteins is assigned a binary label for each of the 32 FunSoCs. This can be visualized as a matrix ***M*** where an entry mij marked as 1 represents that Proteini is annotated as having FunSoCj, or in other words Proteini is positively labelled for FunSoCj. On the contrary, mij marked as 0 means that Proteini does not belong to FunSoCj and is negatively labelled for that FunSoC. Every sequence of the collected set of labelled proteins is positively labeled for at least one FunSoC.

### Dataset Curation and Preprocessing

To build a training and testing dataset for our models, proteins were obtained that were not positively associated with each FunSoC. This was done to avoid tagging every sequence analyzed by SeqScreen with a particular FunSoC. The great majority of biological sequences are benign, so we decided to append the set of curated proteins with a selected set of proteins from SwissProt and labelled them with 0’s for each FunSoC. This forced the model to learn it could neglect assigning FunSoCs to proteins. Further, these proteins were only selected if they had an annotation score greater than 3, to control for the quality of annotation. Once this set of proteins and their respective negative labels were added to the initial list of curated proteins, we extracted relevant features from each of the proteins to be included as features. GO annotations and keywords for each protein were extracted from UniProt. Once extracted, a large binary feature matrix ***F*** was constructed for the total set of proteins. The rows represent each protein in the dataset and the columns represent all possible features of the dataset, (i.e., a union of all the individual features of each protein in the dataset). Each entry fij in the feature matrix ***F***, is a binary value representing presence or absence of a particular featurej for a proteini. Apart from controlling for annotation scores, to further help reduce the effect of noise and non-specific keywords or GO terms from our datasets, we decided to preprocess the feature set to exclude any sparse features that occurred in less than 10 proteins. This reduced the total number of features from over 50k to around 16k features. This was the final feature matrix used for downstream Machine Learning tasks.

### Machine Learning Models

The challenge of assigning FunSoCs to proteins is a multi-class, multi-label classification problem where a given protein can be assigned to any (or none) of 32 different FunSoCs. These are often independent of one another and can be learned individually. Multi-class and multi-label classifications are hard as often these classes have different amounts of training data available. This might make certain labels harder to predict than others and result in a poor classifier that is biased to certain well curated class labels. This also makes accuracy a tricky metric to handle given the imbalance in data labels. From our feature matrix we observed that the number of proteins labelled negatively (i.e., 0) for all FunSoCs greatly outnumbered those with at least one positive label. Though this mirrors the label imbalance in real data, it poses a challenge in learning tasks as the models tend to learn features only from the majority class thereby achieving high accuracy by classifying everything as negative. To address this, we investigate incorporating class weights and sampling techniques into our models. Another challenge often encountered in such tasks is overfitting. By choosing a relatively high number of examples (25% of the training) we carefully monitored the validation and training accuracy to ensure they were similar. We also used regularization techniques such as L1-regularization (Support Vector) and dropout (Neural Networks) to balance weights and reduce overfitting in our models.

Recently, the explainability of predictive models for machine learning has been emphasized in microbiome research[61,62]. To follow this idea of producing explainable results, we used feature selection or two-step modular approaches that aided the interpretability of the models. Though we analyzed 10 models for our FunSoC prediction task, here we describe the top three best-scoring approaches combined with a majority voting scheme. **Fig. 3** illustrates the architectures and parameters of the top three models as part of the ensemble classifier. The first is a two-stage modular pipeline that uses neural networks. For the purposes of this discussion, we describe stage 1 as the detection stage and stage 2 as the classification stage. In the detection stage, we use a multi-layer perceptron with one hidden layer consisting of 200 neurons. The network has a binary output which encodes whether the input sequence is associated with at least one FunSoC. Proteins without FunSoCs are eliminated from downstream classification. Proteins that have at least one FunSoC reach the classification stage which detects FunSoCs associated with a sequence in a multi-label fashion. The architecture of the detection stage consists of one hidden layer with 500 neurons. The output layer contains one neuron per FunSoC that outputs a binary label. For both detection and classification, all internal layers use ReLU activation while the output layers have sigmoid classification. The binary cross-entropy loss function is shown in Eqn. 1. where *y* (0 or 1) is the class label and p is the predicted probability that the observation belongs to class *y*. This is used in conjunction with the Adam optimizer[63] and the models also incorporate a dropout layer with rate 0.2.

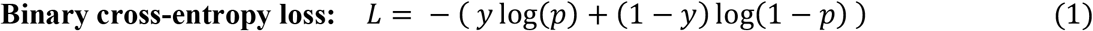

The second model is analogous to the two-stage neural network pipeline except for two major differences. First, the neural networks are replaced with Linear-Support Vector Classifiers (LinearSVC). The LinearSVCs are tuned with training label weights to account for class imbalance and have a binary output for detecting the presence of at least one FunSoC. Second, the classification architecture now consists of different LinearSVCs, one for each FunSoC. Each classification LinearSVC has a binary output indicating the presence of that FunSoC. Both the detection and classification LinearSVCs uses squared hinge loss with L1 penalty (shown in Eq. 2, where *Yi* is the output label, *Xi* is the feature vector of sample *i* and *β* is the vector of weights, *n* is the number of samples and *p* is the number of features), a c-value (*C*) of 0.01 and 4000 iterations for convergence during training.

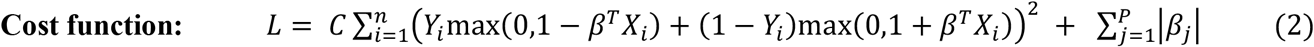

The third best performing model deviates from the two-stage detection and classification pipeline and instead incorporates a feature detection step prior to classification to help with interpretability. The model is a combination of LinearSVCs and neural networks and uses one of each for each FunSoC. In the first step, LinearSVCs are used as a feature selection tool to extract important features for each FunSoC. Since the L1 penalty was used for classification, it assigns a weight of zero to features that are not discriminative towards the FunSoC classification. The LinearSVCs were also augmented with class weights to make the feature selection sensitive to the minority positive labels in each FunSoC. The LinearSVC used an L1 penalty, a c-value of 0.01 and 3000 iterations. Once the features are selected, this new feature set is fed as an input to the neural network for classification. The neural network has one hidden layer with 100 neurons and uses ReLU activation for internal layers and sigmoid activation for the output layer, a dropout layer with rate 0.2 and binary cross-entropy loss. To further lessen the effects of class imbalance, after feature selection random oversampling of the minority class was done prior to training the neural network to balance the number of positive and negative samples in the training set.

The LinearSVCs for all the models were directly incorporated using their scikit-learn[64] implementations. To implement the neural networks, the Keras[65] package was used. Parameter tuning was carried out by varying the c-value (*C*) and testing using different kernels for other non-linear SVCs whereas the number of layers, depth of the neural network, activations, dropout rate, and including class weights was tested for the neural network model. The parameters reported above were consistently the best performing across the parameter space while maintaining a relatively simple architecture and were chosen as the final parameters. The architecture for the three top models is visualized in **Supplementary Figures SF3, SF4** and **SF5**

To combine the strengths of all the classifiers discussed above, we also analyzed an additional model that employed an ensemble majority vote on the outputs of the three models. The ensemble classifier was developed after visualizing performances of the three individual classifiers on hard-to-classify FunSoCs like *develop in host, nonviral invasion, toxin synthase* and *bacterial counter signaling* to try and balance the disparity between precision and recall. To have a model that does not suffer from sub-optimal performances on multiple FunSoCs we reasoned that a majority vote classifier would be a better overarching model for a consistent performance across FunSoCs for downstream applications, especially pathogen detection.

A primary focus during the development of the ML models was to make the feature selection and classification strategies as explainable as possible instead of applying it as “black box” techniques. The interpretability of the models was also imperative for iterative curation where the features and labels could be passed on to the biocurators to potentially curate and refine more examples of proteins belonging to the respective FunSoCs. These refined labels were then fed back into the ML models to obtain the final FunSoC assignments. To minimize variability of our ML results and make SeqScreen analysis more reproducible, ML-based predictions are pre-computed on all of UniProt and is included in the SeqScreen database as a lookup file. This allows users to explicitly view and check the FunSoCs associated with individual UniProt hits and corroborate their biological accuracy.

### Pathogen Sequence Identification

In this work, we provide motivating experiments that underlie an important application of SeqScreen towards pathogen detection. We run SeqScreen on isolate reads obtained from four pairs of well characterized but hard-to-distinguish pathogens namely *E. coli K-12 MG1655 and* pathogenic *E. coli O157:H7*, as well as distinguishing *C. botulinum* from *C. sporogenes*, and *S. dysgalactiae* from *S. pyogenes* in addition to identifying the commensals *S. pyogenes* and *L. gasseri*. To carry out accurate FunSoC annotations, the reads were preprocessed to remove low quality bases and adapters using Trimmomatic[66]. In addition to evaluating SeqScreen, we also ran the set of bacterial reads through Mash dist, Sourmash, PathoScope, Kraken2, KrakenUniq, MetaPhlAn3, and Kaiju. These tools (except PathoScope) were run as part of the MetScale v1.5 pipeline (https://github.com/signaturescience/metscale) using default parameters and a quality trim threshold of 30 with Trimmomatic, k value of 51 with Sourmash, and all other MetScale v1.5 default parameters, tool containers, and databases for analyzing paired-end Illumina reads. We evaluated the results on their respective complete databases as well as a modified version of their database (for Mash dist and PathoScope) in which the entries corresponding to the query genome were removed to simulate a novel or emerging pathogen. In case of *E. coli* the respective strains were removed while in the case of the other bacteria the species (and all strains) were omitted from the database. To facilitate manipulating the Mash database, we created the Mash database from a new version of RefSeq (downloaded November 2020, Release 202). The RefSeq genomes were downloaded using the tool ncbi-genome-download available on conda (https://github.com/kblin/ncbi-genome-download). The genomes downloaded included complete genomes as well as chromosomal sequences *(--assembly-levels complete,chromosome* parameter)

### Sequences from Peripheral Blood Mononuclear Cells in COVID-19 Patients

Sequencing data from three samples of healthy individuals (CRR125445, CRR125456, CRR119890) and three samples of COVID-19 samples (CRR119891, CRR119892, CRR119893) from the study Xiong et al [47]were considered for our analysis. After preprocessing reads through quality control and human read removal (see detailed methods here: https://osf.io/7nrd3/wiki/home/), each sample was passed through SeqScreen v1.2 to obtain the respective set of proteins, FunSoCs, and GO terms outputs. GO terms were parsed with the CoV-IRT-Micro scripts (https://github.com/AstrobioMike/CoV-IRT-Micro), and GO terms were identified that were unique to both the COVID-19 patient samples and viral proteins. The SeqScreen tsv final report was used to connect proteins to GO terms and find all coronavirus reads in the samples. HUMAnN2 was run on the COVID-19 samples to obtain enriched genes and pathways to compare SeqScreen against.

## Availability and requirements

**Project name:** SeqScreen v1.4.11

**Project home page:** https://gitlab.com/treangenlab/seqscreen

**Operating system(s):** Linux

**Programming language(s):** Nextflow, Perl and Python

**Other requirements:** Requirements and Dependencies are listed in the <u>GitLab wiki page</u>. Dependencies can be downloaded by installing SeqScreen via Bioconda: https://anaconda.org/bioconda/seqscreen

**License: GNU GPL V3**

**Restrictions to use by non-academics: None**

## Acknowledgements

Our appreciation goes out to Chris Hulme-Lowe, Danielle LeSassier, Nicolette Albright, Katharina Weber, Veena Palsikar, Oana Lungu, Curt Hewitt, Pravin Muthu, Cynthia DiPaula, Isaac Mayes, Don Bowman, Christopher Grahlmann, and Leslie Parke for their efforts in assisting with the development of the UniProt queries, internal organization of the curation data, and assistance in software development and testing at Signature Science, LLC. We would also like to acknowledge project team contributions of Jason Hauzel, Kristófer Thorláksson, Manoj Deshpande, Brendan Joyce, Garrit Nickel, Steffen Matheis, Larissa Wagnerberger, Vijayadhaarani Vijayaarunachalam, Mikael Lindvall, and Adam Porter at Fraunhofer CMA USA for their support in software quality assurance and development of the HTML report generator, Jeremy Selengut of the University of Maryland for sharing his insights into viral pathogenesis, Jim Gibson of Signature Science, LLC for graphics development, and Letao Qi, Jacob Lu, and Chris Jermaine of Rice University for insightful discussions and work specific to the machine learning algorithms. Thanks to Jody Proescher, Ron Jacak, Kristina Zudock and Briana Vecchio-Pagan from Johns Hopkins University Applied Physics Lab for their helpful discussions about pathogenesis ontologies and assistance with deployment and testing of our software on their servers. SeqScreen software development has significantly benefited from the feedback of Elizabeth Vitalis and Ian Fiddes at Inscripta and Cory Bernhards’ team at U.S. Army Combat Capabilities Development Command Chemical Biological Center. We are thankful for all the time and effort provided be end users to test early versions of our software, recommend improvements, and guide its application to current challenges in pathogen detection, synthetic biology, and genome engineering. Finally, we would like to thank IARPA and all our SeqScreen end users for their helpful feedback and support over the course of the Fun GCAT program.

## Ethics approval and consent to participate

Not applicable

## Consent for publication

Not applicable

## Author Contributions

K.T. and T.T. designed the SeqScreen concept. A.B., B.K., D.A, T.T., M.D., L.E., Z.Q., and D.N. developed the software. G.G, A.K., and K.T. designed and implemented the biocuration framework. A.B., B.K. T.T. and S.S. designed and implemented the machine learning framework. N.S. and M.P. designed outlier detection. A.B., B.K., Z.Q., E.R produced results for benchmarking. A.B., L.E., G.G., S.S., N.S., M.P, K.T., and T.T. contributed to writing the manuscript. All authors read and approved the manuscript.

