## Supplementary Material for "SeqScreen: Accurate and Sensitive Functional Screening of Pathogenic Sequences via Ensemble Learning"

### SUPPLEMENTARY DATA

#### **SD1.** Database construction

PFam, NCBI nucleotide, and MEGARes databases were downloaded from their locations and are compiled into indexed databases using default parameters for their respective tools. The REBASE restriction enzyme database was downloaded and converted to a FASTA file to be used with MUMmer. The Uniref100 FASTA file was downloaded as well as the ID mapping from the UniProt FTP server. GO-terms, PATRIC IDs, and STRING IDs were all mapped to their corresponding UniProt IDs in the Uniref100 FASTA file. This way, when sequence hits are identified in SeqScreen, there is no need to perform a lookup of the hit's UniProt ID to obtain the metadata, as the necessary metadata is now part of the sequence header in the fasta file. It is worth noting that this resulted in a larger storage requirement for the DIAMOND and BLAST databases. In addition to stitching in the UniProt metadata, all UniParc ID entries were removed to remove sequences with low annotation qualities. BLASTX and DIAMOND databases were then constructed from the resulting pruned UniRef100 FASTA file with default parameters. The FunSoC database was created from the UniProt (release-2019_05), downloaded on 07/10/2019. UniParc IDs were filtered from the database, and a final set of 30275173 proteins were used as the prediction set for the majority vote ML model.

#### **SD2.** GO term and pathway analysis in COVID-19 samples.

The GO terms analysis for the COVID-19 study can be found here: https://osf.io/8j4d2 and the HUMAnN2 analysis can be found here: https://osf.io/y5mzk/

#### **SD3.** Benchmarking

To determine the ability of SeqScreen to accurately characterize functional and taxonomic labels on short genomic segments, we benchmarked it against some of the top-performing functional annotation and taxonomic classification tools. As many of the current functional annotation tools are not tailormade for annotating short sequences or subsequences of proteins with Gene Ontology terms, we analyzed the results of the tools over a range of different lengths, including over the entire length of the protein sequence to provide a holistic estimate of their performance. On taxonomic datasets, we compared the performance of the tools on both the genus and species taxonomy levels. We used seven different metagenomic datasets with unambiguous taxonomic ground truths closely resembling commonly found environmental metagenomes. Similar datasets have also been used in previous taxonomic benchmarking studies[54]. In the following sections, we describe the datasets, experimental outline and tools considered for our benchmarking experiments.

##### **SD3.1** Functional Benchmarking

Annotating short genomic or protein sequences with functional annotation terms is a challenging task for current tools. Unlike full length proteins, these sequences can have a very specific set of GO terms associated with it which may be distinct from full sequences. Also, shorter sequences are inadequately characterized in full length protein databases and often hit multiple proteins which can confound any annotation analysis. Here, we aim to measure the accuracy of each tool to assign annotation labels to proteins and observe how changes in the length of the sequence affects their ability. To compare different tools on the accuracy of assigning functional annotation related Gene Ontology (GO) terms we use datasets from previous iteration of the Critical Assessment of Functional Annotation (CAFA 3) which is a benchmarking challenge to test computational approaches and tools on their ability to assign GO terms in three different categories: Molecular Function and Biological Processes. In each iteration of the CAFA challenge, a batch of training and testing sets of proteins are released. The training sets consist of proteins with previously established and experimentally verified labels and are used by the tools to "learn" sequence features that correspond to specific set(s) of GO terms. The testing sets are typically hard-to-classify proteins, a subset of which are experimentally annotated during the challenge and are the proteins the tools are scored upon. For the purposes of our work, we chose training dataset from the CAFA 3 challenge for the functional annotation benchmarking task. Further, since we envision SeqScreen to be applicable in settings where functional characterization of the sequences is of primary importance, we only benchmark against Molecular Function and Biological Process GO term annotations.

##### **SD3.1.1** Dataset Preparation

To create the benchmarking dataset tuned to our application, we considered the initial training set from the CAFA 3 challenge and pruned it to select a subset of proteins that have been annotated by at least one of the FunSoCs by our team of biocurators. In total, we selected 250 such proteins across 32 FunSoCs from the CAFA training sets. We chose the CAFA training set since it had CAFA provided labels for each protein to benchmark against. To simulate proteins of multiple lengths from the full-length proteins in the training set, we randomly selected a starting point on the protein and extend it to contiguous sequences of length 34, 50, 67 and 80 amino acids in addition to the full protein sequence. Thus, we obtained 250 sequences for each of the four lengths and the corresponding full-length sequences. We then transferred the Molecular Function and Biological Process GO terms for the full sequence as the truth labels for all the shorter sequences. As SeqScreen only accepts DNA sequences as input, we used the EMBOSS Backtranseq tool[S1] to get the corresponding DNA sequences for our protein set.

##### **SD3.1.2** Functional Annotation Tools

We considered some of the top performing tools from previous CAFA challenges for our benchmarking analysis. Since SeqScreen is an open-source installable command line software, we did not consider any web-server based tools even if they were among the top CAFA performers (e.g., NetGo) and only considered tools which contained or relied on publicly downloadable databases, open source dependencies, and could be run on a command line. We narrowed the list of tools to PANNZER2, DeepGOPlus and eggNOG-mapper. Here, we briefly describe the salient features of each of the tools and analyze the diverse set of approaches they use to solve the GO term annotation challenge. The tool eggNOG-mapper relies on fast orthology based lookups to assignments based on precomputed clusters and phylogenies that are transferred on to the query sequence based on the hits. Depending on the database used, eggNOG-mapper can use either DIAMOND or HMMER 3 to identify seed orthologs. It then discards distant ortholog matches and transfers functional descriptors based on one-to-one or all closest ortholog hits depending on the user options. PANNZER2 uses a Sequence Similarity Result List (*SSRL*) obtained after a database search which is partitioned into clusters based on a term frequency-inverse document frequency *(tf-idf)* based Description Similarity Measure (DSM). A specialized regression model is then used to perform an enrichment analysis of GO terms specific to each of the clusters. PANNZER2 has performed consistently as one of the top performing tools in previous iterations of CAFA. Finally, we will discuss DeepGOPlus, one of the top-performing deep learning-based methods for functional annotation of proteins. It is based on a multi-filter convolutional neural network that learns sequence level features *de novo*. Proteins are encoded using a one-hot encoding scheme and filters of different sizes are used to capture patterns of residues at different granularities. These filters are then combined using individual maxpools and passed to a fully connected feed-forward network that outputs labels for each GO class. DeepGOPlus shows comparable performance to the top performing tools in the most recent CAFA 3 challenge, and the authors claim that DeepGOPlus would have been amongst the top three best predictors for predicting Cellular Component terms and the second-best performing method for Biological Process and Molecular Function GO term evaluations in the CAFA 3 challenge[S2].

##### **SD3.1.3** Functional Annotation Results

Though advances have been made to label protein sequences with corresponding GO terms, this task remains a challenging problem especially for shorter sequences. To assess the ability of SeqScreen to accurately annotate proteins with GO terms and short sequences, we analyzed the performance of SeqScreen on 250 proteins from the CAFA 3 training datasets. Additionally, all the proteins chosen for benchmarking had associated FunSoC labels and hence were sequences containing markers considered to be harmful to humans by a panel of subject matter experts to make the benchmarking relevant for our study. In addition to full-length sequences, we also looked at protein fragments of four different lengths (34, 50, 67 and 80 amino acids) sampled from the same set of 250 proteins. For the purposes of this benchmarking experiment, we compared against three other functional annotation tools shown to do well in previous CAFA competitions and other benchmarking studies, namely PANNZER2, eggNOG-mapper and DeepGOPlus. It is important to note that these tools were developed to annotate unknown full-length protein sequences, and, to our knowledge, no other leading tools have been developed to assign GO terms to short protein subsequences. From the CAFA study we only considered those tools that are available to be run as command line interfaces, like SeqScreen, and hence omit top performing purely web-based servers in that competition (ex: NetGO, INGA 2.0). Both DeepGOPlus and PANNZER2 also outputted confidence values along with GO terms. We set the lower threshold of confidence values to consider as 0.1 after a sweep of all values to determine the one with the best performance. For the full-length sequences, **Fig. SDF1** illustrates the results obtained on this dataset. SeqScreen had the highest precision among all the tools considered in the benchmarking study. We also observed that, although DeepGOPlus and PANNZER2 had lower precision than SeqScreen, they had higher recall. The performance of eggNOG-mapper on complete sequences was slightly better than on subsequences.

For the partial sequences (34 aa, 50 aa, 67 aa, and 80 aa) sampled from full-length proteins, we evaluated SeqScreen's performance versus the CAFA 3 top performer. Over all four sub-lengths of the proteins, SeqScreen had the highest F1 score (**Fig. SDF2**). For the 34 amino acid length case, both DeepGOPlus and eggNOG-mapper had poor precision (0.76 and 0.38 respectively) compared to the rest of the tools. PANNZER2 had the highest recall of all the tools and the second best F1 score behind SeqScreen. All tools besides eggNOG-mapper demonstrated better recall than precision. As we increased the length to 50 amino acids, eggNOG-mapper showed the most improvement in terms of precision and recall, increasing from 0.38 to 0.69 and 0.11 to 0.40 respectively, though it still had lower accuracy than the other three tools. DeepGOPlus also showed an improvement in precision from 0.76 to 0.82. At length 67 amino acids, PANNZER2's recall remained the best among the tools whereas its precision dropped from 0.93 to 0.92. Also, DeepGOPlus's precision starts to pick up, increasing from 0.82 to 0.91. Moving from 67 to 80 amino acids, a major change was observed as DeepGOPlus attained a higher precision than PANNZER2 (0.95 compared to 0.91) and marginally surpassed PANNZER's F1 score (0.952 compared to 0.945). The recall of all the tools increased at 80 amino acids length compared to 67 amino acids.

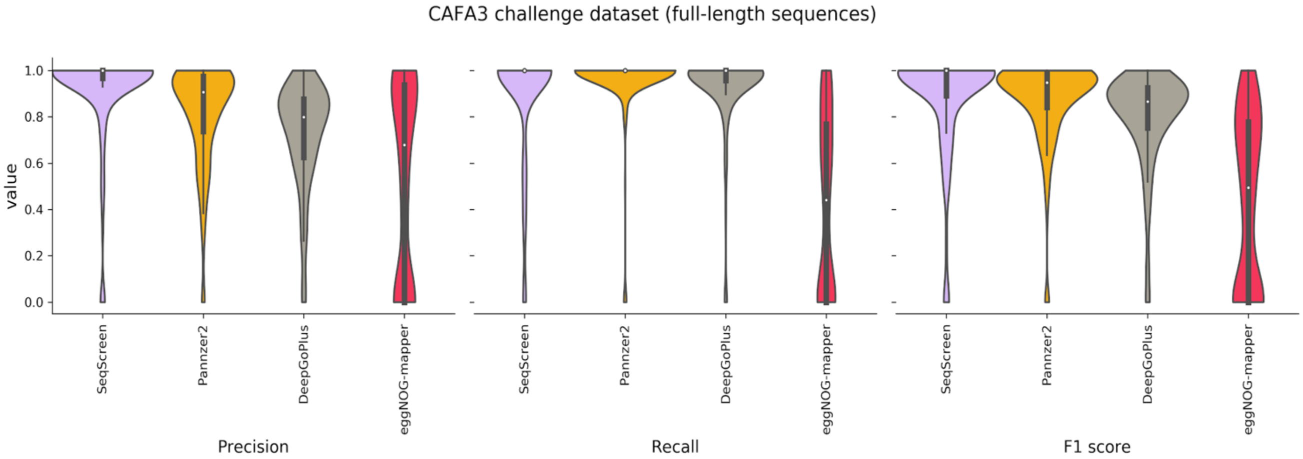


**Fig. SDF1. Functional annotation results on full-length proteins.** SeqScreen shows the highest precision of all tools followed by PANNZER2 and DeepGOPlus. eggNOG-mapper's precision is the lowest among all tools. In terms of recall, PANNZER2 followed by DeepGOPlus has the highest performance. They are followed by SeqScreen and eggNOG-mapper. SeqScreen still reported the highest F1 score among all tools.

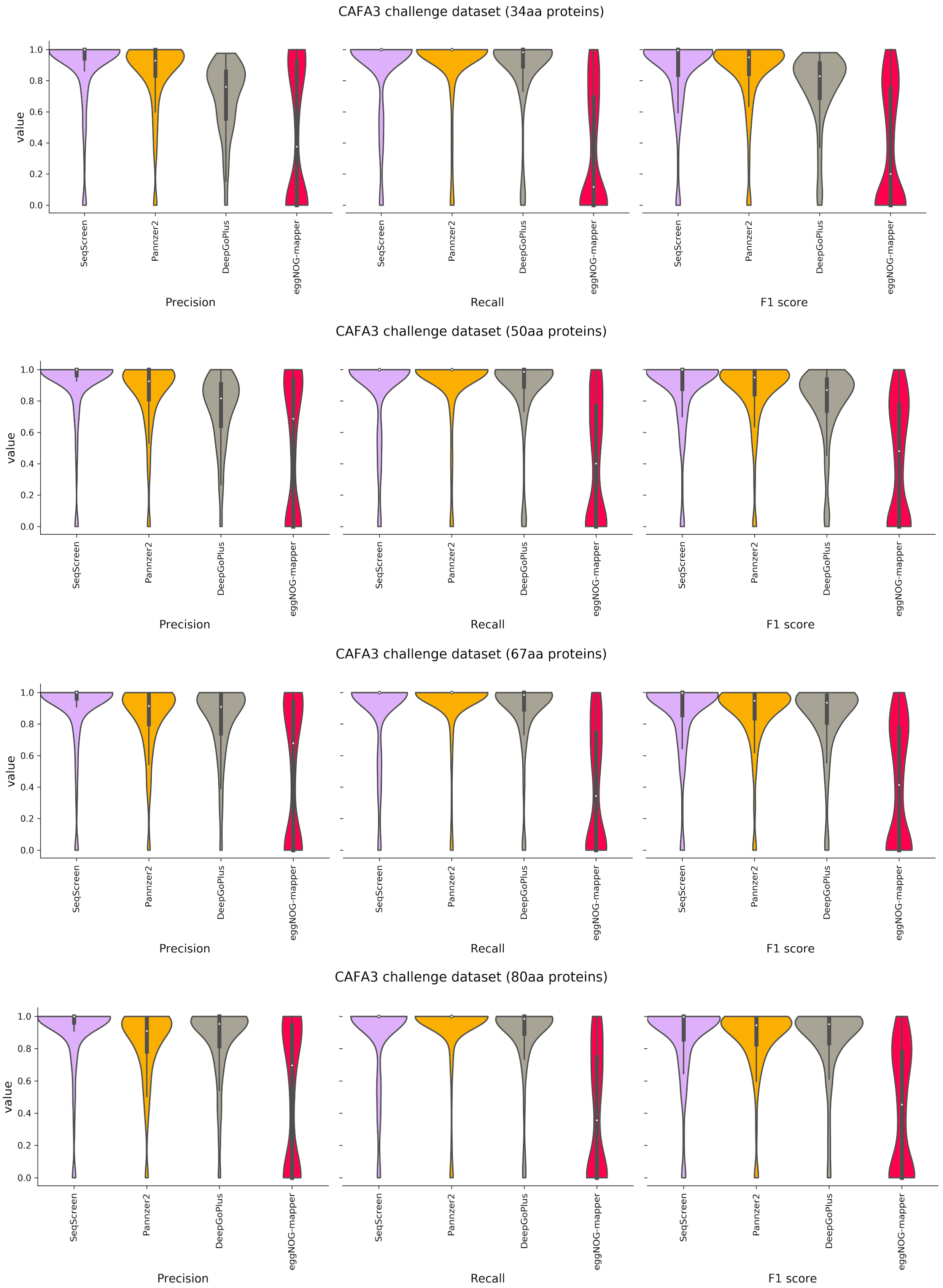


**Fig. SDF2. Functional annotation results on 250 proteins obtained from CAFA-3 training datasets for 34, 50, 67 and 80 amino acids (aa).** The first figure on each panel represents precision followed by recall and F1 score. We observe that SeqScreen consistently achieves the best performance for both precision and recall and by extension the best F1 score for all the sub-lengths. The order of the tools is in decreasing order of their median F1 score. PANNZER2 maintains the highest recall through all the lengths while its precision increases slightly from 34aa to 67aa. eggNOG-mapper increases its precision, recall and F1 score from 34aa to 67aa but only sees marginal improvement thereafter. DeepGOPlus improves its precision above that of PANNZER2 at 80aa.

#### **SD3.2** Taxonomic Benchmarking

Accurate and sensitive taxonomic classification is imperative to the task of characterizing genomic sequences. Recently there have been enormous advances made in k-mer-based taxonomic classifier space in terms of time and memory efficiency, but accurate species level labelling remains a challenge. From a pathogen detection standpoint, there are additional confounding factors separating extremely close taxa. Detection is even more challenging and crucial if one of the closely related candidate organisms is extremely pathogenic and only differs because of the presence of a toxin gene (e.g., the *BotA* gene in *Clostridium botulinum* vs. *Clostridium sporogenes*). Misclassification of such closely related organisms is commonly observed when classifying short DNA segments using the prevalent k-mer-based methods. These methods rely on a predefined k-mer-length for the query and match them to the curated taxonomic databases specific to a given tool and are not tuned towards identifying markers of pathogenicity in sequences. Here, we compared SeqScreen to popular k-mer-based metagenomic taxonomic classifiers using seven different simulated datasets. These datasets have a well-defined unambiguous truth set that can be used to compare the sensitivity and specificity of the classifications at different taxonomic levels. For the purposes of our benchmarking, we analyzed classification by all tools at the species and genus levels.

##### **SD3.2.1** Dataset Preparation

We considered seven simulated datasets developed by Rachid Ounit used in previous tool benchmarking and comparison studies[S3]. These were modeled on distinct microbial habitats based on studies that characterized real metagenomes found in the human body (e.g., buccal, gut) and in the natural or built environment (e.g., city parks/medians, houses, soil). Another dataset was created to represent the organisms of the New York City subway system as described in the study of Afshinnekoo et al.[S4], using the same methodology. Briefly, 100-bp Illumina reads were simulated using the ART simulator[S5] with default error and quality base profiles from sets of reference sequences at 30X coverage and post-processed to remove ambiguously mapped reads at the species levels using MEGAN[S6]. Hence, the reads thus obtained map unambiguously to a single species within the NCBI/RefSeq database and can be easily scored to compare classification tools.

##### **SD3.2.2** Taxonomic Classification Tools

We considered six different metagenomic classifiers for our analysis: Kraken[S7], Kraken 2[S8], KrakenUniq[S9], MetaOthello[S10], Kaiju[S11] and Centrifuge[S12]. As introduced earlier, based on our initial in-house testing we found that Centrifuge was a top performer in several test datasets and its advantages over traditional k-mer based methods (at a marginal runtime cost) was instrumental in it being included as part of a merged taxonomic identification module in SeqScreen. Centrifuge was also benchmarked as a standalone tool in order to obtain a fair comparison to SeqScreen.

We now briefly describe the internal sequence analysis algorithm for each method. Kraken uses an efficient search database that contains Least Common Ancestor (LCA) - k-mer pairs indexed by minimizers for fast lookups. Kraken computes root to leaf (RTL) paths for each taxon level hit and assigns the lowest taxon in the most weighted RTL path to the sequences. In case of ambiguity, it tends to assign the LCA taxon to the sequence. Kraken2 is an update to Kraken with modifications to make database lookups faster and reduce its memory footprint while maintaining similar accuracy levels. Kraken2 uses a probabilistic hashmap to store minimizer-LCA pairs and does away with storing all k-mers. In addition, the hashmap is designed to have improved memory access locality and thread scaling.

KrakenUniq is similar to Kraken except that it also maintains a separate HyperLogLog data structure to estimate k-mer cardinalities and trades higher precision and recall at the cost of extra memory. MetaOthello uses a probabilistic hashing of taxon specific k-mer signatures. It also maintains a novel data structure called I-Othello which holds a pair of hash tables mapping k-mers to taxids. Lastly, Kaiju is one of the few tools that incorporates protein sequence information for taxonomic assignment by carrying out a six-frame translation similar to SeqScreen. It also employs a Burrows-Wheeler Transform[S13] (BWT) based compression of the protein database to facilitate efficient query search and retrieval.

##### **SD3.2.3** Taxonomic Benchmarking Results

Having shown that SeqScreen provides state-of-the-art functional characterization of short genomic sequences, we now illustrate its ability to accurately classify sequences at various taxonomic levels. To that effect, we compared SeqScreen's taxonomic identification pipeline with six popular metagenomic taxonomic classifiers on seven different datasets modeled on a real metagenomic dataset. Fig. SD1. Shows that the combined taxonomic identification pipeline in SeqScreen performed better than other tools on the benchmarking metagenomic datasets at both the species and genus levels.

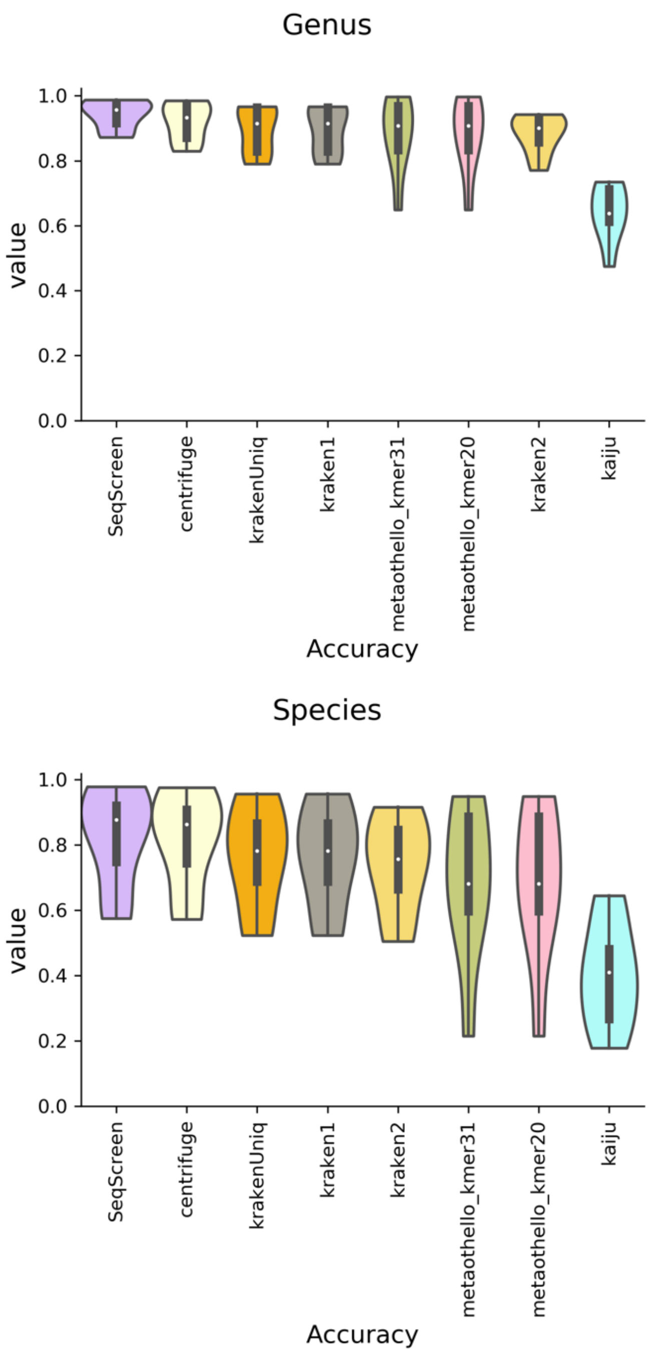


**Fig. SDF3. Genus (left panel) and species (right panel) level accuracy for taxonomic benchmarking**. Accuracy was calculated per sequence and averaged over the entirety of all datasets. SeqScreen showed the highest accuracy at both species and genus level. The combined pipeline (Centrifuge + DIAMOND) in SeqScreen performs marginally better than centrifuge. All of the Kraken tools perform very similar with KrakenUniq performing the best of the three. Kraken is followed by MetaOthello and Kaiju. We observe that the k-mer size in MetaOthello does not contribute to the variability of its output on our datasets.

From the results it was also apparent that all the classifiers performed much better at the genus level than at the species level. SeqScreen's accuracy increased from 0.9 in species classification to upwards of 0.95 at the genus level. This was closely followed by Centrifuge. KrakenUniq and Kraken2 had the same accuracy at species (0.78) and genus (0.9) levels. Kraken2 had better accuracy than MetaOthello at species level but does poorly when compared to it at genus level. We also observed that k-mer size has no effect on MetaOthello's ability to classify sequences on this benchmarking dataset. Overall, SeqScreen had higher accuracy than the current competing tools over seven distinct metagenomic benchmarking datasets.

#### **SD4.** Example of UniProt queries

UniProt queries used for FunSoCs: https://rice.app.box.com/s/3ahojrtpoqj5vcdbocd1tac1rrn5ocow

#### **SD5.** Current FunSoC database

Current FunSoC DB: https://obj.umiacs.umd.edu/seqscreen/funsocs_20.9.26.tsv

#### Supplementary Data References


S1. Madeira, Fábio, et al. "The EMBL-EBI search and sequence analysis tools APIs in 2019." *Nucleic acids research* 47.W1 (2019): W636-W641.

S2. Kulmanov, Maxat, and Robert Hoehndorf. "DeepGOPlus: improved protein function prediction from sequence." *Bioinformatics* 36.2 (2020): 422-429.

S3.Ounit, R. & Lonardi, S. Higher classification sensitivity of short metagenomic reads with CLARK-S. *Bioinformatics* **32**, 3823–3825 (2016).

S4.Afshinnekoo, E. *et al.* Geospatial Resolution of Human and Bacterial Diversity with City-Scale Metagenomics. *Cell Syst* **1**, 97–97.e3 (2015).

S5.Huang, W., Li, L., Myers, J. R. & Marth, G. T. ART: a next-generation sequencing read simulator. *Bioinformatics* vol. 28 593–594 (2012).

S6.Huson, D. H., Auch, A. F., Qi, J. & Schuster, S. C. MEGAN analysis of metagenomic data. *Genome Res.* **17**, 377–386 (2007).

S7.Kraken, C. C. *Benezit Dictionary of Artists* (2011) doi:10.1093/benz/9780199773787.article.b00101018.

S8.Wood, D. E., Lu, J. & Langmead, B. Improved metagenomic analysis with Kraken 2. *Genome Biol.* **20**, 257 (2019).

S9.Breitwieser, F. P., Baker, D. N. & Salzberg, S. L. KrakenUniq: confident and fast metagenomics classification using unique k-mer counts. *Genome Biol.* **19**, 198 (2018).

S10.Liu, X. *et al.* A novel data structure to support ultra-fast taxonomic classification of metagenomic sequences with k-mer signatures. *Bioinformatics* **34**, 171–178 (2018).

S11.Menzel, P., Ng, K. L. & Krogh, A. Fast and sensitive taxonomic classification for metagenomics with Kaiju. *Nat. Commun.* **7**, 11257 (2016).

S12. Kim D, Song L, Breitwieser F, Salzberg S; Centrifuge: Rapid and sensitive classification of metagenomic sequences. *Genome Research* (2016) 26(12) 1721-1729

S13. Burrows, M. & Wheeler, D. J. *A Block-sorting Lossless Data Compression Algorithm*. (1994).

#### SUPPLEMENTARY TABLES

**Supplementary Table ST1**. Overview of the 32 FunSoCs. FunSoCs are listed along with their definitions, an example of a protein with te FunSoC, and if they can be found in viruses (V), bacteria (B), or eukaryotes (E). FunSoCs are not found in archaea, so that is not included in this table**.**

| **FunSoC Name** | **Category** | | | **FunSoC Definition** | **Protein Example** |
|---|---|---|---|---|---|
| | **V** | **B** | **E** | | |
| Disable organ | X | X | X | Disables an organ, but not necessarily by killing individual cells. This includes neurotoxins that block receptor conductance, superantigens, and toxins that degrade lung function, gut function, or are involved in the disruption of blood homeostasis. | *Clostridium botulinum* Botulinum neurotoxin type G (Q60393) |
| Cytotoxicity | X | X | X | Kills cells by inhibiting a vital process such as translation, directly lysing the cells through pore formation, or destabilizing the plasma membrane | *Escherichia coli* Hemolysin E (Q68S89) |
| Degrade ecm | X | X | X | Mediates the loosening of host tissue connectivity including reducing cell-to-cell adherence and cell-to-substrate adherence as well as proteolytic destruction of the host extracellular matrix (ECM) | *Trimeresurus gramineus* Zinc metalloproteinase/disintegrin (P0C6E8) |

| | | | | | |
|---|---|---|---|---|---|
| Induce inflammation | X | X | X | Directly activate host inflammatory pathways to cause damage | *Salmonella typhimurium* Guanine nucleotide exchange factor SopE (O52623) |
| Bacterial counter signaling | | X | | Bacterial suppression of host immune signaling within host cells to avoid inflammatory responses | *Mycobacterium tuberculosis* Protein LppM (O53505) |
| Viral counter signaling | X | | | Viral suppression of host immune signaling within host cells to avoid inflammatory responses | *Epstein-Barr virus* Tegument protein BLRF2 (P03197) |
| Resist complement | X | X | X | Provide resistance to host complement components | *Candida albicans* pH-regulated antigen PRA1 (P87020) |
| Counter immunoglobulin | X | X | X | Binds, sequesters, destroys, or otherwise neutralizes host immunoglobulin | *Staphylococcus aureus* Immunoglobulin G-binding protein A (P02976) |
| Plant RNA silencing | X | | | Counters host plant defenses by triggering the degradation of specific sequences, pre-translation | *Turnip yellow mosaic virus* 69 kDa protein (P20131) |
| Resist oxidative | | X | | Enable resistance of host oxidative killing | *Vibrio cholerae* Peptide methionine sulfoxide reductase MsrA/MsrB (Q9KLX6) |
| Suppress detection | X | | | Viral interference with host immune components involved in processing or display of parasite antigens on the surface of host cells | *Human cytomegalovirus* Unique short US6 glycoprotein (P14334) |
| Avirulence plant | X | X | X | Virulence effector proteins of plant pathogens that typically suppress immune signaling of the host plant | *Pseudomonas syringae* Effector protein HopAB2 (Q01723) |
| Host gtpase | | X | | Target host small GTPases | *Legionella pneumophila* Phosphocholine transferase AnkX (Q5ZXN6) |
| Host transcription | X | X | X | Target host transcription to inhibit or activate | *Influenza A virus* RNA-directed RNA polymerase catalytic subunit (Q809M3) |
| Host translation | X | X | X | Target host translation, usually to inhibit | *Human adenovirus* Shutoff protein (P11824) |
| Host ubiquitin | X | X | X | Target host ubiquitination machinery | *Human SARS coronavirus* Replicase polyprotein 1a (P0C6U8) |
| Host xenophagy | X | X | | Target host xenophagy/autophagy | *Rotavirus A* Non-structural glycoprotein 4 (Q82028) |
| Nonviral invasion | | X | X | Mediates bacterial or eukaryotic pathogen invasion into host cells | *Plasmodium falciparum* Cysteine-rich protective antigen (Q8IFM8) |
| Viral invasion | X | | | Mediates viral invasion into host cell | *Bean golden yellow mosaic virus* Capsid protein (P0CK34) |
| Viral movement | X | | | Enable movement of viral parasite within a host cell or tissue | *Squash leaf curl virus* Nuclear shuttle protein (P21935) |
| Virulence activity | X | X | X | Inclusive of a wide variety of virulence activities, including those that did not fit under another FunSoC category | *Phytophthora infestans* RxLR effector protein Avr3a (E2DWQ7) |
| Host Cell cycle | X | X | | Target host cell cycle component or induce alterations in the host cell cycle | *Cowpox virus* Ankyrin repeat domain-containing protein CP77 (P12932) |
| Host Cell death | X | X | X | Target host apoptotic cell death pathways either to inhibit or activate | *Human papillomavirus* Protein E6 (P03126) |
| Host cytoskeleton | X | X | | Target host cytoskeletal components or induce alterations in the host cytoskeleton | *Pseudomonas aeruginosa* Putative aldolase class 2 protein PA3430 (Q9HYH5) |

| | | | | | |
|---|---|---|---|---|---|
| Secreted effector | | X | | Secreted bacterial effectors of unknown function | *Salmonella choleraesuis* Secreted effector protein PipB2 (Q57KZ6) |
| Antibiotic resistance | | X | X | Counters the effect of antibiotics administered to inhibit the growth or vital functioning of bacterial or eukaryotic parasites. Note: These are not specific to pathogens. | *Bacillus cereus* β-lactamase (P01424) |
| Develop in host | | X | X | Involved in the growth or development of a bacterial or eukaryotic parasite within the host | *Mycobacterium tuberculosis* Putative diacyglycerol O-acyltransferase Rv2285 (P9WKB5) |
| Nonviral adhesion | | X | X | Mediates adherence of nonviral parasites to host cells | *Mycobacterium tuberculosis* 60 kDa chaperonin 2 (P9WPE7) |
| secretion | | X | | Bacterial secretion system components (T1SS - T8SS), including chaperones | *Burkholderia mallei* Secretion apparatus protein BsaZ (A3MCG6) |
| Toxin synthase | | X | X | Enzymes that synthesize or modify toxins, particularly mycotoxins | *Penicillium expansum* ABC transporter patM (B6RAL1) |
| Viral adhesion | X | | | Mediates viral adherence to host cells | *Vaccinia virus* Cell surface-binding protein (O57211) |
| Virulence regulator | | X | X | Bacteria or eukaryotic parasites transcription factor of virulence | *Magnaporthe oryzae* C2H2 finger domain transcription factor CON7 (G4N5Q2) |

**Supplementary Table ST2:** Performance of other machine learning models on the FunSoC dataset. Mean Positive label (PL) precision and recall denote the average precision and recall on testing samples associated with a given FunSoC. Numbers in bold indicate the highest scores for Micro-F1, PL precision and recall. Majority voting ensemble is an ensemble classifier of models

4,6 and 11 each of which scored highest on the three metrics Micro-F1, PL precision and recall. Abbreviations: NN – neural networks, OS - oversampling

| Model | Accuracy | Exact Match Ratio | Micro- F1 score | Mean Positive label Precision | Mean Positive l[illegible] Recall |
|---|---|---|---|---|---|
| 1. Linear SVCs | 99.97% | 99.29% | 93.49% | 74.19% | 62.02% |
| 2.SVC + NN (OS) | 99.61% | 90.16% | 49.08% | 55.86% | 85.39% |
| 3.Weighted Joint NN | 99.64% | 95.64% | 49.03% | 34.06% | 80.49% |
| 4.TS NN | 99.97% | 99.24% | 93.59% | **88.93%** | 69.88% |
| 5.TS SVC | 99.97% | 99.23% | 91.52% | 67.99% | 57.95% |
| 6.Bl. SVC + NN (OS) | 99.97% | 99.24% | **98.59%** | 87.59% | 81.80% |
| 7.Chi2 + NN (OS) | 99.97% | 99.25% | 93.18% | 84.01% | 71.60% |
| 8.ExtraTree + NN (OS) | 99.97% | 99.37% | 94.96% | 85.11% | 73.60% |
| 9. AdaBoost + NN (OS) | 99.97% | 99.38% | 94.08% | 87.76% | 85.22% |
| 10.TS Bl.NN | 99.97% | 99.38% | 93.83% | 80.46% | 67.41% |
| 11. TS Bl.SVC | 99.96% | 98.93% | 86.92% | 73.82% | **88.69%** |
| 12. Majority Voting Ensemble (4+6+11) | 99.97% | 99.34% | 99.24% | **90.03%** | 82.73% |

**Supplementary Table ST3.** Expanded table containing the full list of genes and FunSoCs for isolate genomes reported by SeqScreen. Abbreviated gene names are listed if at least one read from the gene had an UniProt e-value < 0.0001, was assigned a FunSoC, and was from the expected genus (i.e., Escherichia or Shigella, Clostridium, Streptococcus, Lactobacillus). FunSoCs with at least one gene that met the criteria for detection in at least one of the isolates were included in the table. The removal of genes from genera that were not expected in these bacterial isolates allowed for removal of genes that were likely derived from likely contaminating organisms (e.g., PhiX Illumina sequencing control). After genes passed the thresholds for detection, the total number of reads that SeqScreen assigned to each gene was recorded and listed in parentheses after the gene abbreviation in this table. Red cells with bold font indicate genes with FunSoCs that most strongly differentiated the pathogenic potential of the two bacterial isolates being compared. The results compared include two different E. coli strains, one commensal (*E. coli K12 MG1655*) and one pathogen (*E. coli O157:H7*), two different Clostridium species, including one rare pathogen (*C. sporogenes*) and one harmful toxin-producing pathogen (*C. botulinum*), two different Streptococcus pathogens from different Streptococcus Groups (*S. pyogenes* and *S. dysgalactiae*), and two different commensal species.

| Isolate (accession #) | disable organ | cytotoxicity | degrade ecm | induce inflammation | secreted effector | secretion | antibiotic resistance | bacterial counter signaling | counter immuno-globulin | virulence regulator |
|---|---|---|---|---|---|---|---|---|---|---|
| ***E. coli K12 MG1655***<br>(DRR198806) | | *hlyE* (22) | | | *tir* (1) | *gspD* (1,776)<br>*gspE* (1,380)<br>*gspL* (1,148)<br>*gspF* (1,123)<br>*yghF* (1,097)<br>*yghE* (820)<br>*gspJ* (573)<br>*gspH* (489)<br>*gspG* (423)<br>*gspI* (398)<br>*gspC* (338)<br>*tatA2* (326)<br>*tatE* (33)<br>*lsrA* (5)<br>*GSP* (4)<br>*hofF* (2)<br>*escN* (1) | *pbpA* (2,505)<br>*pbpB* (2,404)<br>*mdtF* (2,285)<br>*mdtO* (2,110)<br>*arnA* (1,611)<br>*mdtP* (1,498)<br>*mdtQ* (1,382)<br>*mdtG* (1,279)<br>*dacB* (1,245)<br>*emrB* (1,239)<br>*macA* (1,228)<br>*mdtE* (1,167)<br>*mdtL* (1,138)<br>*fsr* (1,130)<br>*bla* (1,052)<br>*emrA* (1,011)<br>*mdtN* (1,009)<br>*mdfA* (1,009)<br>*uppP* (881)<br>*arnD* (864)<br>*arnB* (814)<br>*arnC* (795)<br>*mdtH* (713)<br>*mdtM* (661)<br>*tehB* (436)<br>*lpxD* (248)<br>*marB* (170)<br>*acrA* (164)<br>*acrZ* (154)<br>*tehA* (151)<br>*pmrD* (82)<br>*eptA* (36)<br>*nimT* (28)<br>*rcbA* (27)<br>*ralR* (17) | | | |
| ***E. coli O157:H7***<br>(DRR198804) | | ***stxB* (195)**<br>*hlyE* (83) | | | *espF(U)* (1,054)<br>*tir* (761) | *etpE (2,241)*<br>*pulD-like* (2,209)<br>*hlyB* (2,008)<br>*etpF* (1,390)<br>*etpJ* (950)<br>*escN* (942)<br>*W0021* (909)<br>*etpH* (823)<br>*etpD* (776)<br>*etpG* (654)<br>*etpC* (454) | *mdtF* (1,997)<br>*arnA* (1,366)<br>*pbpB* (1,349)<br>*mdtO* (1,295)<br>*pbpA* (1,176)<br>*mdtP* (959)<br>*mdtQ* (898)<br>*mdtG* (877)<br>*bla* (856)<br>*mdtM* (853)<br>*emrB* (797)<br>*mdtL* (786) | | | *espF(U)* (1,054)<br>*nleF* (347) |

| | | | | | | | | | | |
|---|---|---|---|---|---|---|---|---|---|---|
| | | | | | | *tatA2* (236)<br>*sntF* (230)<br>*lsrA* (1) | *mdtE* (775)<br>*dacB* (773)<br>*macA* (764)<br>*arnB* (744)<br>*fsr* (712)<br>*mdfA* (634)<br>*mdtN* (632)<br>*arnD* (619)<br>*uppP* (576)<br>*mdtH* (568)<br>*arnC* (485)<br>*emrA* (293)<br>*acrA* (127)<br>*acrZ* (124)<br>*tehB* (98)<br>*rcbA* (43)<br>*ralR* (34)<br>*eptA* (28)<br>*tehA* (3)<br>*lpxD* (2)<br>*nimT* (2)<br>*marB* (1) | | | |
| ***C. sporogenes***<br>(SRR8758382) | | | | | | *T3SA* (1) | *bla* (1,042)<br>*uppP* (660)<br>*pbpA* (137) | *moaC* (129) | | |
| ***C. botulinum***<br>(SRR8981313) | ***botA* (193)** | *orf-X2* (4) | ***botA* (177)** | | | | *pbpA* (317)<br>*bla* (216)<br>*uppP* (205) | *moaC* (61) | | ***botA* (177)** |
| ***S. dysgalactiae***<br>(SRR12825903) | | *SLO* (8,076) | | | | *comYC* (1,669) | *uppP* (3,897)<br>*pilA* (9) | | ***spg* (5,241)** | |
| ***S. pyogenes***<br>(ERR1735064) | | *SLO* (1,552) | | ***speB* (1,008)**<br>***speH* (325)** | | *comYC* (293) | *uppP* (619) | | | |
| ***S. salivarius***<br>(SRR11910125) | | | | | | *cglC* (1)<br>*comGC* (295) | *mprF* (1,347)<br>*uppP* (811)<br>*pbpA* (8) | | | |
| ***L. gasseri***<br>(SRR11910134) | | | | | | | *mprF* (244)<br>*aac(3)* (138) | | | |

#### Gene Names

- *aac(3)* = Aminoglycoside N(3)-acetyltransferase
- *acrA* = Multidrug efflux pump subunit AcrA
- *acrZ* = Multidrug efflux pump accessory protein AcrZ
- *arnA* = Bifunctional polymyxin resistance protein ArnA, Bifunctional polymyxin resistance protein arnA domain protein, Putative bifunctional polymyxin resistance protein ArnA, Polymyxin resistance protein ArnA_DH, UDP-glucuronic acid decarboxylase / Polymyxin resistance protein ArnA_FT, UDP-4-amino-4-deoxy-L-arabinose formylase
- *arnB* = UDP-4-amino-4-deoxy-L-arabinose--oxoglutarate aminotransferase
- *arnC* = Undecaprenyl-phosphate 4-deoxy-4-formamido-L-arabinose transferase
- *arnD* = Probable 4-deoxy-4-formamido-L-arabinose-phosphoundecaprenol deformylase ArnD
- *bla* = Beta-lactamase, Beta-lactamase (Fragment)
- *botA* = Botulinum neurotoxin type A2, Botulinum neurotoxin type A, Botulinum neurotoxin, Botulinum neurotoxin A5, Bont/A1, and BoNT/A3
- *cglC* = Competence protein CglC
- *comGC* = Competence protein ComGC
- *comYC* = Competence protein, Putative competence protein
- *dacB* = D-alanyl-D-alanine carboxypeptidase DacB
- *emrA* = Multidrug export protein EmrA

- *emrB* = Multidrug export protein EmrB
- *eptA* = Phosphoethanolamine transferase EptA
- *escN* = Type III secretion system, ATPase LEE associated, EscN/YscN/HrcN family type III secretion system ATPase, Putative EscN
- *espF(U)* = Secreted effector protein EspF(U)
- *etpC* = EtpC protein
- *etpD* = Type II secretion protein
- *etpE* = Type II secretion system protein E
- *etpF* = EtpF protein
- *etpG* = EtpG protein
- *etpH* = EtpH protein
- *etpJ* = EtpJ protein
- *fsr* = Fosmidomycin resistance protein
- *GSP* = General secretion pathway protein, Putative general secretion pathway for protein export (GSP)
- *gspC* = General secretion pathway protein C, General secretion pathway protein GspC (Fragment), Hypothetical type II secretion protein GspC, Putative type II secretion system protein C
- *gspD* = General secretion pathway protein D, Putative secretin GspD, Type II secretion system protein D
- *gspE* = Putative type II secretion system protein E, Type II secretion system protein E
- *gspF* = General secretion pathway protein F, Putative type II secretion system protein F, Type II secretion system protein F
- *gspG* = General secretion pathway protein G, Type II secretion system core protein G
- *gspH* = General secretion pathway protein H, General secretion pathway protein H (Fragment), General secretion pathway protein GspH, Type II secretion system protein H
- *gspI* = General secretion pathway protein I, Putative type II secretion system protein I
- *gspJ* = General secretion pathway protein J, Type II secretion system protein J, General secretion pathway protein GspJ
- *gspL* = General secretion pathway protein L, General secretion pathway protein L (Fragment), Putative type II secretion system protein L, Type II secretion system protein L
- *hlyB* = Alpha-hemolysin translocation ATP-binding protein HlyB, Hemolysin B
- *hlyE* = Hemolysin E
- *lpxD* = UDP-3-O-(3-hydroxymyristoyl)glucosamine N-acyltransferase
- *lsrA* = Autoinducer 2 import ATP-binding protein LsrA
- *macA* = Macrolide export protein MacA
- *marB* = Multiple antibiotic resistance protein MarB
- *mdfA* = Multidrug transporter MdfA
- *mdtE* = Multidrug resistance protein MdtE
- *mdtF* = Multidrug resistance protein MdtF
- *mdtG* = Multidrug resistance protein MdtG
- *mdtH* = Multidrug resistance protein MdtH
- *mdtL* = Multidrug resistance protein MdtL
- *mdtM* = Multidrug resistance protein MdtM
- *mdtN* = Multidrug resistance protein MdtN
- *mdtO* = Multidrug resistance protein MdtO
- *mdtP* = Multidrug resistance outer membrane protein MdtP
- *mdtQ* = Multidrug resistance outer membrane protein MdtQ, Putative multidrug resistance outer membrane protein MdtQ
- *moaC* = Cyclic pyranopterin monophosphate synthase
- *mprF* = Phosphatidylglycerol lysyltransferase
- *nimT* = 2-nitroimidazole transporter
- *nleF* = Effector protein NleF
- *orf-X2* = Neurotoxin accessory protein
- *pbpA* = Penicillin-binding protein 1A
- *pbpB* = Penicillin-binding protein 1B

- *pilA* = Prepilin-type cleavage/methylation N-terminal domain protein
- *pmrD* = Signal transduction protein PmrD
- *pulD-like* = PulD-like protein
- *ralR* = Endodeoxyribonuclease toxin RalR
- *rcbA* = Double-strand break reduction protein
- *SLO* = Streptolysin O, Thiol-activated cytolysin
- *sntF* = Putative ATP-dependent Clp protease
- *speB* = Streptopain
- *speH* = Exotoxin type H
- *spg* = Immunoglobulin G-binding protein G
- *stxB* = Shiga toxin subunit B
- *tatA2* = Sec-independent protein translocase protein TatA
- *tatE* = Probable Sec-independent protein translocase protein TatE
- *tehA* = Tellurite resistance protein TehA
- *tehB* = Tellurite methyltransferase
- *tir* = Translocated intimin receptor Tir
- *uppP* = Undecaprenyl diphosphate, Undecaprenyl-diphosphatase, Undecaprenyl-diphosphatase 1, Undecaprenyl-diphosphate phosphatase, Undecaprenyl-diphosphate phosphatase (Fragment)
- *W0021* = W0021
- *yghE* = Putative type II secretion system L-type protein YghE
- *yghF* = Putative type II secretion system C-type protein YghF, Secretion pathway, C-type protein

**Supplementary Table ST4.** Software and commands used for Functional and Taxonomic Benchmarking

| SOFTWARE | VERSION | DATABASE | COMMAND |
|---|---|---|---|
| PANNZER2 | 2.0 | UniProt | python runsanspanz.py -R --PANZ_FILTER_PERMISSIVE --PANZ_MINLALI 5 -o <output file> -i <input file> |
| DeepGOPlus | 1.0.0 | UniProt | ./predict.sh <input file> <output file> |
| eggNOG-mapper | 2.0.1 | CoG, eggNOG Protein DB | python2 emapper.py --hmm_evalue 0.0001 --hmm_score 15 -i <input file> --output <output file> -m diamond - -cpu 10 |
| DIAMOND | 0.9.26 | UniRef 100 (Curated) | diamond blastx -q <fasta>\ <br> -d <database> \ <br> -o <out>\ <br> --evalue <evalue>\ <br> --threads <threads >\ <br> --block-size 200 \ <br> --index-chunks 1 \ <br> --salltitles \ <br> --more-sensitive \ <br> --min-orf 10 \ <br> --masking 0 \ <br> --top 5 \ <br> -f 10 |
| Centrifuge | 1.0.4 | RefSeq (archaeal, bacterial, viral) | centrifuge -x <centrifuge_path> + <centrifuge_index > -f -U <input name> -S, <output file> --report-file <report file name> -p 15 |

| | | | |
|---|---|---|---|
| Kraken | 1.0 | Kraken Standard DB (RefSeq) | kraken -db <kraken database> --threads 10 --output <output file> --fasta-input <input file> |
| Kraken2 | 2.0.8-beta | Kraken2 Standard DB (RefSeq) | kraken2 --db <kraken2 database> --threads 10 --report <report file> --use-mpa-style --output <output file> |
| KrakenUniq | 0.5.8 | Kraken Standard DB (RefSeq) | krakenuniq --db <database> --threads 10 --report-file <report file> --fasta-input <input file> |
| metaOthello | (last update - 2018) | RefSeq | ./classifier \ bacterial_{20/31}mer_L12.index \ <output directory> \. 20/ 31 \ 10 \ fa \ SE \ bacterial_speciesId2taxoInfo.txt \ NCBI_names_file.txt\ <input file> \ |
| Kaiju | 1.7.2 | NCBI nr | kaiju -t <look up table> -f <reference input file> -i <input file> -o <output file> -z 80 |

**SUPPLEMENTARY FIGURES**

**Supplementary Figure SF1.** SeqScreen's taxonomic workflow.

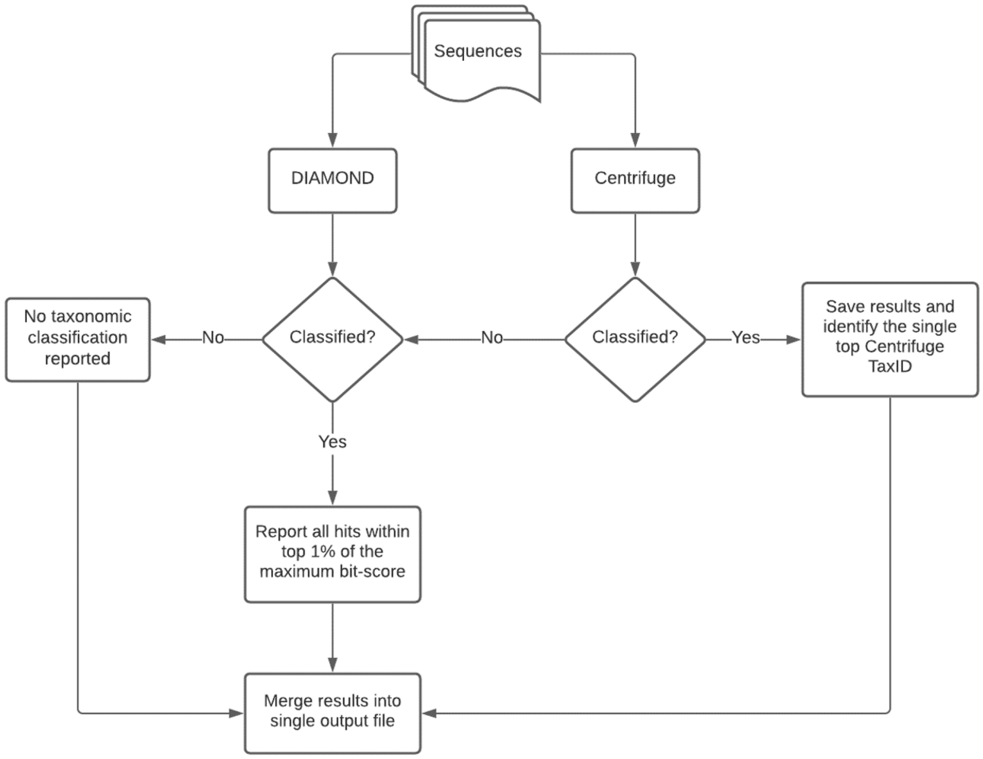

**Supplementary Figure SF2:** GO-term visualization. Within the SeqScreen HTML report, there is an interactive visualization of GO terms available for each sequence queried, and an example of that is shown here.

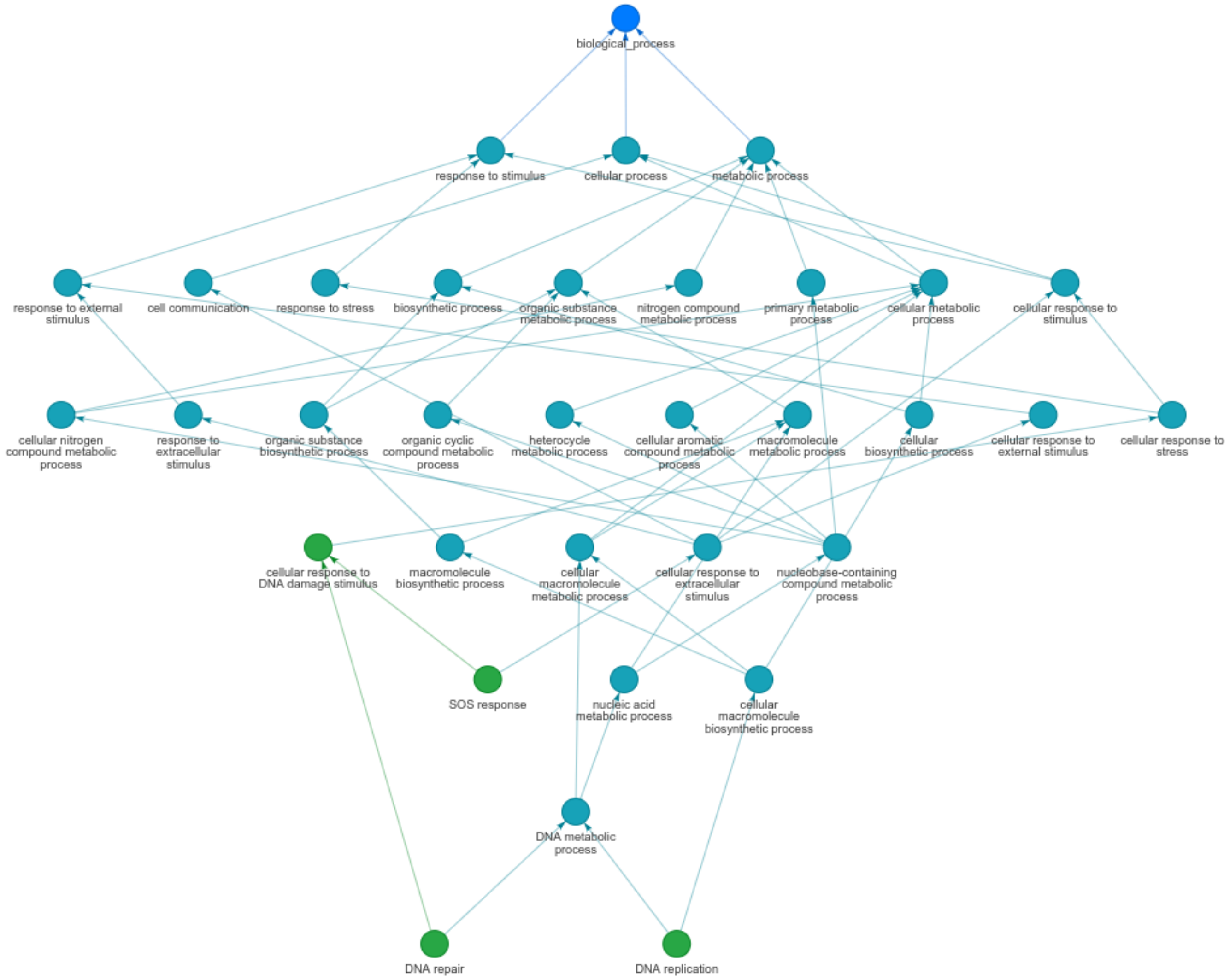
